# *In Silico* studies provide new structural insights into *trans*-dimerization of β_1_ and β_2_ subunits of the Na^+^,K^+^-ATPase

**DOI:** 10.1101/2024.08.13.607722

**Authors:** Gema Ramírez-Salinas, Jorge L. Rosas-Trigueros, Christian Sosa Huerta, Liora Shoshani, Marlet Martínez-Archundia

## Abstract

The Na^+^,K^+^-ATPase is an electrogenic transmembrane pump located in the plasma membrane of all animal cells. It is a dimeric protein composed of α and β subunits and has a third regulatory subunit (γ) belonging to the FXYD family . This pump plays a key role in maintaining low concentration of sodium and high concentration of potassium intracellularly. The α subunit is the catalytic one while the β subunit is important for the occlusion of the K^+^ ions and plays an essential role in trafficking of the functional αβ complex of Na^+^,K^+^-ATPase to the plasma membrane. Interestingly, the β_1_ and β_2_ (AMOG) isoforms of the β subunit, function as cell adhesion molecules in epithelial cells and astrocytes, respectively. Early experiments suggested a heterotypic adhesion for the β_2_. Recently, we reported a homotypic trans-interaction between β_2_-subunits expressed in CHO cells. In this work we use *In Silico* methods to analyze the physicochemical properties of the putative homophilic trans-dimer of β_2_ subunits and provide insights about the *trans*-dimerization interface stability. Our structural analysis predicts a molecular recognition mechanism of a *trans*-dimeric β_2_-β_2_ subunit and permits designing experiments that will shed light upon possible homophilic interactions of β_2_ subunits in the nervous system.

**Author summary:** The adhesion molecule on glia (AMOG) is the β_2_ isoform of the β-subunit of the Na^+^-pump that is localized in the nervous system, specifically in astrocytes. It was shown that it mediates Neuron-Astrocyte interaction, promoting neurite outgrowth and migration during brain development. In recent years we have shown that the ubiquitous β_1_ isoform is a homophilic adhesion molecule in epithelia and therefore we hypothesized that β_2_ could also interact as a homophilic adhesion protein. In a previous work we show that fibroblasts (CHO) transfected with the human β_2_ subunit of the Na^+^-pump become adhesive. Moreover, protein-protein interaction assay in a co-culture of cells transfected with β_2_ tagged with two different markers (His_6_ and YFP) reveal a positive interaction between the β_2_-subunits. In the present work, we apply bioinformatics methods to analyze and discuss the formation of a *trans*-dimer of β_2_-subunits. Our *In Silico* study predicts a relatively stable dimer with an interface that involves the participation of four out of the seven N-glycosylation sites. Nevertheless, interacting interface and the dynamics of the β_2_-β_2_ *trans*-dimer is different from that of the β_1_-β_1_ dimer; it involves different surfaces and therefore it explains why β-subunits can not form mixed (β_1_-β_2_) *trans*-dimers.

## 1. INTRODUCTION

The Na^+^, K^+^-ATPase, a ubiquitous plasma-membrane ion pump plays a crucial physiological role in all animal cells. Indeed, the resultant ion and electrochemical gradients are essential for many physiological processes and in the brain, about 50% of the ATP is consumed by the Na^+^,K^+^-ATPase [1]. Na^+^,K^+^-ATPase is a P-type ATPase, an oligomeric enzyme that consists of three subunits: α, β and *γ* [2,3]. This work is focused on the β-subunit.

The Na^+^,K^+^-ATPase β subunit is part of the functional core of the pump and is required for its trafficking to the plasma membrane. Mammals express three β subunit isoforms, β_1_, β_2_ and β_3_. It has a small intracellular, N-terminal domain (30 amino acids), a single transmembrane helix, and a large extracellular, C-terminal domain of about 240 amino acids [4,5]. The different β isoforms have distinct tissue and cell-type specific expression profiles [6,7]. There are three conserved disulfide bonds in the extracellular domain, which are important for forming a stable pump [8], and the extracellular domain has three, eight, and two glycosylation sites in β_1_, β_2_, and β_3_, respectively [9,10]. Functionally, β_2_ has the strongest effects on the kinetic properties of the pump, reducing the apparent potassium affinity and raising the extracellular sodium affinity compared to β_1_ and β_3_ [11]. The different β isoforms and the variation in their post-translational modifications facilitate regulated Na^+^, K^+^-ATPase activity, adapted to different tissues and to environmental changes.

The β subunit is important for the occlusion of the K^+^ ions and plays an essential role in trafficking of the functional αβ complex of Na^+^, K^+^-ATPase to the plasma membrane [12]. Apart from the role of β subunit in regulating the pump activity, a role in cell-cell adhesion has been also proposed [13].

With this regards, [13] have suggested that the Na^+^,K^+^-ATPase acts as a cell adhesion molecule by binding to the Na^+^,K^+^-ATPase molecule of a neighboring cell by means of *trans*-dimerization of their β_1_ subunits. Following, it was demonstrated that a direct homotypic interaction between β_1_-subunits of neighboring cells, takes place between polarized epithelial cells [14,15] identified the amino acid region crucial for the species-specificity of this trans-interaction [16] completed the description of the adhesion interface between the extracellular-domains of the dog β_1_-subunits.

Earlier, the group of Schachner identified an adhesion molecule on glia (AMOG) that functions as a neural recognition molecule mediating neuron-glia interactions that promotes migration and neurite outgrowth [17,18]. This adhesion molecule was later identified as the β_2_-subunit of the Na^+^,K^+^-ATPase and was named β_2_/AMOG [19]. Their works suggested a heterophilic interaction between AMOG and an unknown molecule at the neuron membrane [19,20].

The crystal structure analysis of the Na^+^,K^+^-ATPase β_1_ subunit in the E2 state as published by Shinoda and colleagues, marked a significant milestone by revealing the atomic structure of the extracellular domain of the β_1_ subunit [PDB: 2ZXE [21]. Notably, the extracellular C-terminal domain of the protein adopts an Ig-like β-sheet sandwich configuration, consistent with *in silico* predictions [22]. Intriguingly, although many adhesion and non-adhesion proteins feature domains with an immunoglobulin-like (Ig-like) topology, structural alignments of the β_1_-subunit extracellular domain against well-studied cell adhesion molecules do not reveal any structural homologs to β subunits. Upon detailed examination, three distinctive features of the β subunit family members emerge: 1. The Ig-like fold with a unique topology, interrupted by a long α-helix secondary structure. 2. An atypical β-sheet disposition in relation to classical Ig folds. 3. The β subunit fold contains extensive loops, resulting in a length twice that of a typical Ig domain. Furthermore, the structural relationship between the β_1_ subunit and the catalytic α subunit suggests that the C-terminal fold must exhibit greater rigidity compared to the typical flexibility seen in adhesion domains, such as cadherin-domains [23]. Further works including mutational analysis combined with *In Silico* studies have identified the residues at the dog β_1_ surface that participate in β_1_-β_1_ interaction [15,16].

Although It is well accepted that both isoforms β_1_ and β_2_ function as adhesion molecules in epithelia and in the nervous system, respectively there is almost no information regarding the adhesion mechanism of β_2_/AMOG isoform.

Very recently it has been published that β_2_ acts as an homophilic adhesion molecule when expressed in CHO fibroblasts and MDCK epithelial cells [24]. Cell-cell aggregation, protein-protein interaction assays as well as *In Silico* studies were carried out to confirm cell-cell adhesion mediated by β_2_-β_2_ *trans*-interaction. With these results the authors localized the putative interacting surface in a docked model and suggested that the glycosylated extracellular domain of β_2_/AMOG, can make an energetically stable trans-interacting homodimer.

In the present work we have built homotypic dimers of the human β_1_ and β_2_ subunits by employing protein-protein docking analysis, and submitted them to molecular dynamics simulations (MDS) which provide detailed information about their dimeric conformation and specific differences in their interfaces. We also investigated the role of the glycosylation in the interface stabilization of the Human β_1_ and β_2_ dimeric complexes.

## 2. RESULTS

### 2.1 Molecular modeling of human ATP1B1 and ATP1B2

The β-subunit of the sodium pump is a membrane protein with a single transmembrane helix and most of the mass folded as a Ig-like β-sandwich at the extracellular space [16,22]. Since the structure of the extracellular domain is stable and active [14,16,25,26,27], we decided to analyze its adhesive properties without the cytoplasmic and transmembrane domains as they do not participate in the β-β trans-interaction.

The three dimensional (3D) model of the extracellular domain of human Na^+^,K^+^-ATPase β_1_ subunit (*ATP1B1*) was built by considering the crystal structure of the Na^+^,K^+^-ATPase 3WGU from wild boar (*Sus scrofa*) and the Fasta Sequence of Uniprot (P05026). In Figure 1A the 3D model of the extracellular domain of β_1_ subunit is depicted. The three N-glycosylation sites: Asn158, Asn193 and Asn265 at the surface of the extracellular domain, the three disulphide bridges and the characteristic Ig-like β-sandwich structure are shown. Validation of the 3D model was carried out by employing Ramachandran plots, where it could be seen that 99% of the residues are included in permitted zones of the protein (Figure 1B).

**Figure 1.**
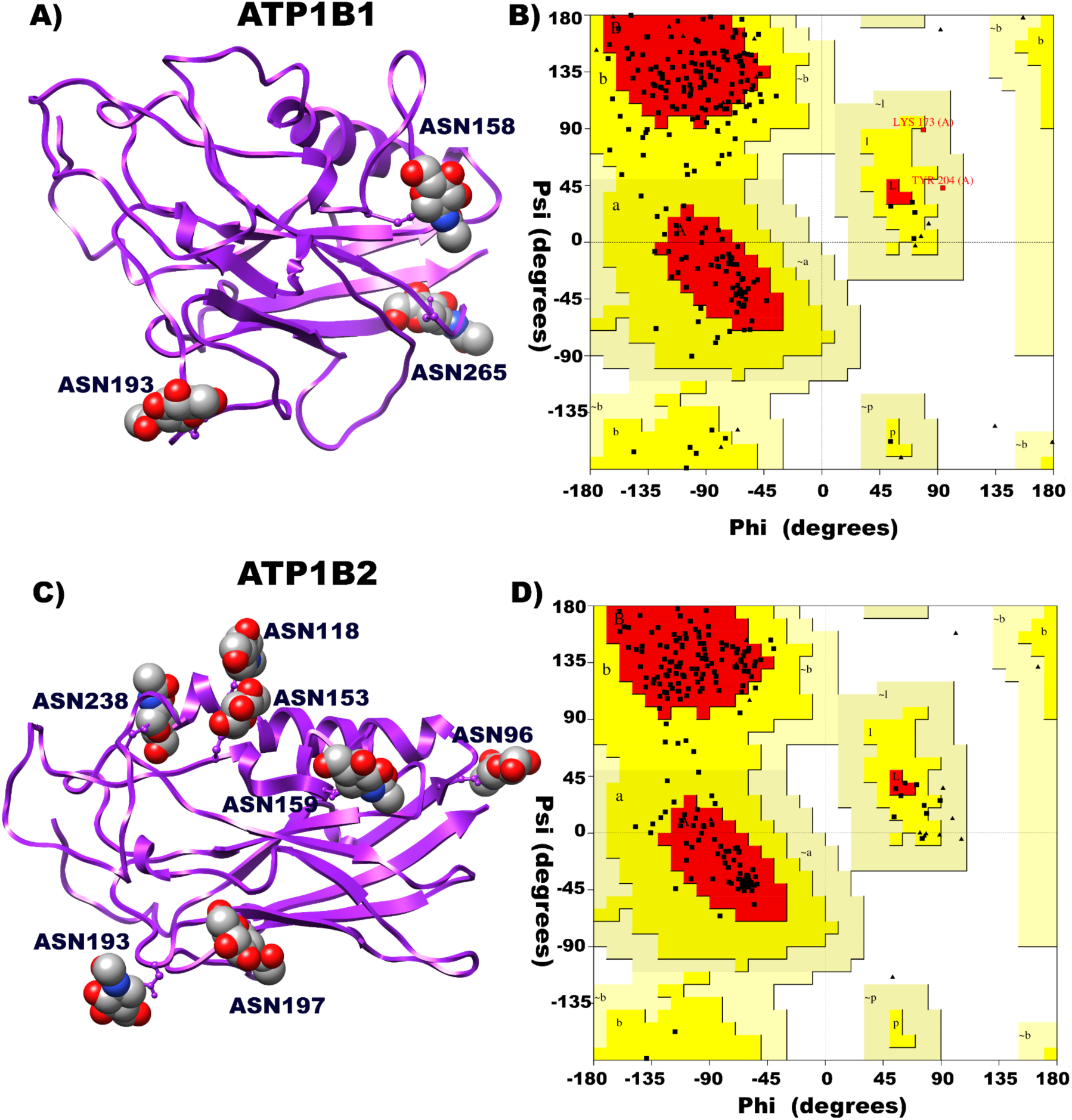
Three-dimensional structure of monomeric β_1_ subunit (*ATP1B1*) and β_2_ subunit (*ATP1B2*). A) 3D structure of the β_1_ subunit monomer including its glycosylations. B) Ramachandran plot of the β_1_ subunit monomer where it can be seen that none of the residues of the proteins are included in the disallowed regions. C) 3D structure of the β_2_ subunit monomer including its glycosylations. D) Ramachandran plot of the β_2_ subunit monomer where it can be seen that none of the residues of the proteins are included in the disallowed regions.

Since no crystal structure was available for the β_2_ subunit of any species, the 3D model of the extracellular domain of human ATP1B2 was built by considering the crystal structure of the homologous (Identity: 40% and Convergence: 98%) pig gastric H^+^,K^+^-ATPase- 5YLU and the Fasta Sequence of Uniprot (P14415). In Figure 1C, the following structural features of the extracellular domain of β_2_ subunit are shown: seven N-glycosylation sites, three disulphide bridges and a characteristic Ig-like β-sandwich structure. Validation of the 3D models was carried out by employing Ramachandran plots, where it could be seen that 100% of the residues are included in permitted zones of the protein (Figure 1D).

### 2.2 Building of the dimers β_1_-β_1_ and β_2_-β_2_

The molecular docking of the extracellular domains of both β_1_-β_1_ and β_2_-β_2_ subunits was performed by using HDOCK Server. In that protein-protein docking process, the most energetically favorable conformers were chosen, for β1-β1 that of -193.04 kcal/mol and for β2-β2 that of -274.99 kcal/mol) by means of the HDOCK Server. Among the favorable conformers, as a second structural criteria, we considered only the trans-dimers for further analyses. In the present work we considered pertinent to include the glycosylations in modeling, docking and MD simulations since it was demonstrated that N-glycosylation of both extracellular domains of β_1_ and β_2_ subunits are crucial for cell-cell adhesion [16,24]. In Figure 2 the selected trans-dimers were depicted.

**Figure 2.**
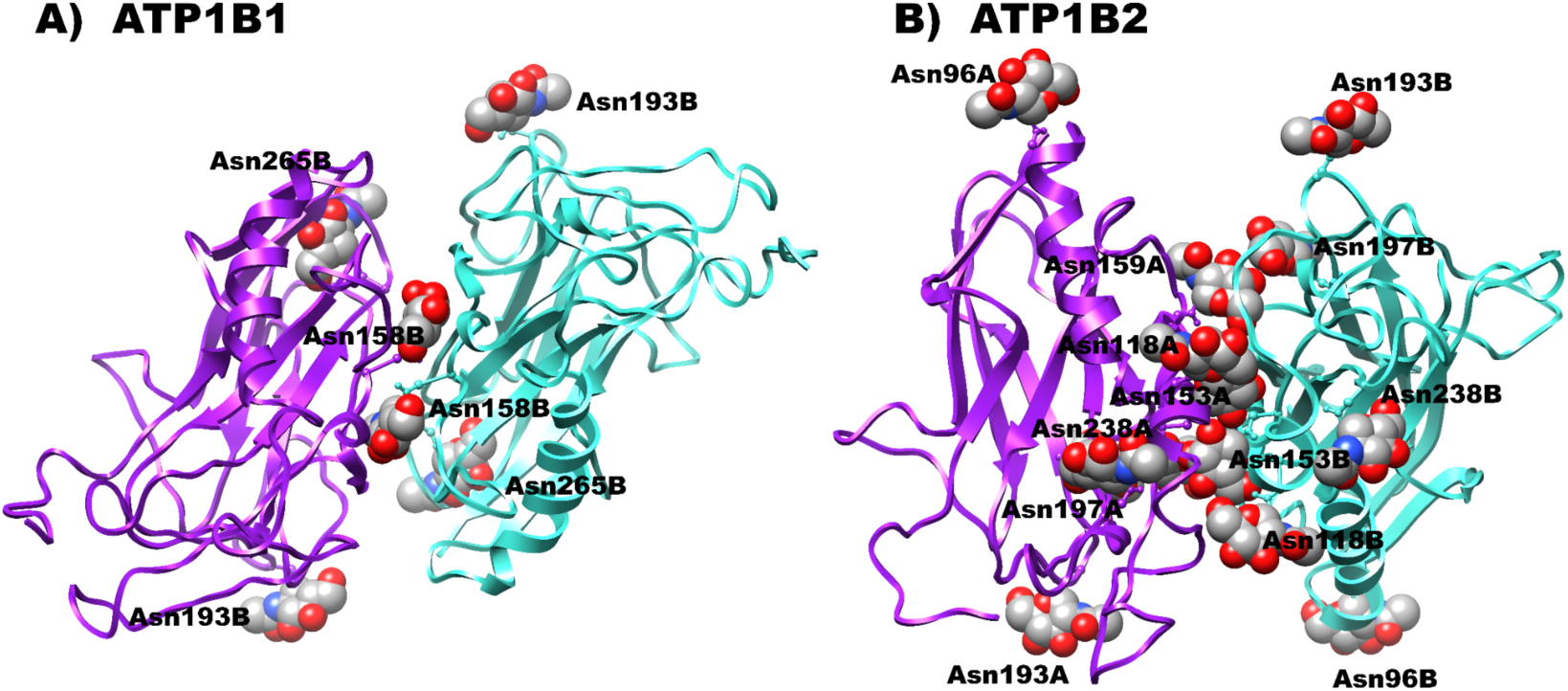
Dimeric 3D structure of β_1_ subunit and β_2_ subunit in *trans* orientation. A) Dimeric structure of β_1_-β_1_. B) Dimeric structure of β_2_-β_2_. For both cases Chain A is colored in purple and Chain B is colored in blue. Glycosylations are marked in balls and sticks.

### 2.3 Molecular Dynamics Simulations of the dimers β_1_-β_1_ and β_2_-β_2_

Molecular dynamics simulations (MDS) were carried out on both dimeric complexes depicted in Figure 2, and trajectories were run for 200 ns. Furthermore, structural analysis was done with the Carma Program as described in “Methods”. In agreement with previous results with dog *ATP1B1* [16], the RMSD values for the soluble ectodomain of human β_1_-β_1_are within the range of 6-8Å eventhogh, the present model includes the three glycosylated residues.

From the structural analysis of the dimers (Figure 3) we could not see remarkable structural differences between β_1_-β_1_ and β_2_-β_2_ dimers. Nevertheless, the surface residues that constitute the adhesion interface in the two dimers were different. Therefore, we decided to analyze both interfaces to get a better understanding about their formation and stability.

**Figure 3:**
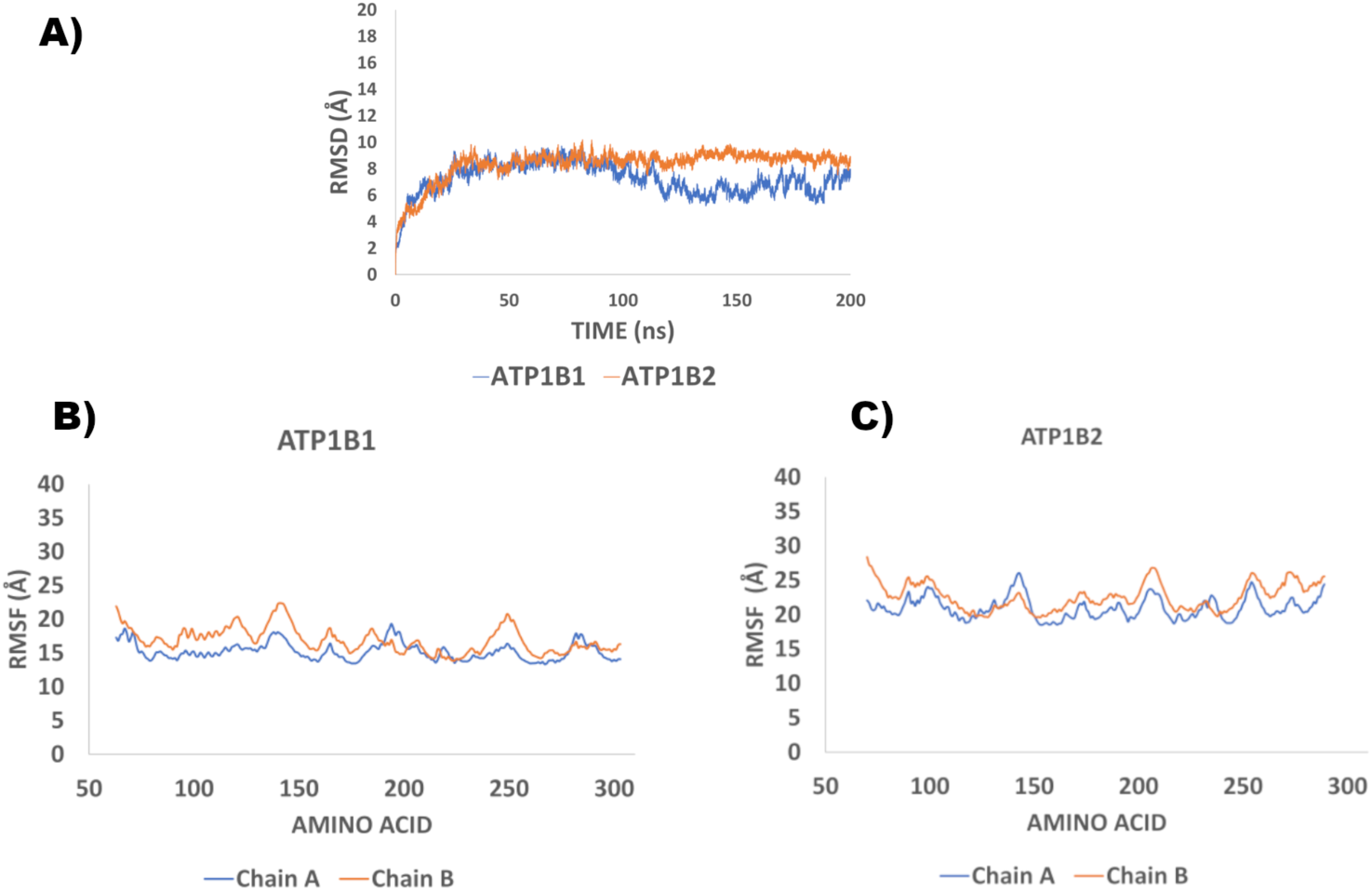
Structural Analysis of ATP1B1 and ATP1B2 dimers: A) Root mean square deviation analysis (RMSD) of the dimers β_1_ and β_2_ . Root mean square fluctuation (RMSF) analysis of the alpha carbons of the dimers β_1_-β_1_ (B) and β_2_-β_2_ (C).

### 2.4 Analysis of interactions in the β-β interfaces

Protein-protein interactions are important for normal biological processes since they play a key role in the regulation of cellular functions that affect gene expression and function [28]. In this work we present an analysis of the residues at the interface of protein-protein interaction, thus providing information about the stability and specificity of the complex. In the analyses of the interfaces, the properties to be considered include: hydrogen bonding, buried surface area and hydrophobicity among others [29]. PPCHEK server was employed to get an insight on the non-bonded interactions that are present in the dimeric complexes (β-β) obtained from the MD simulations. These interactions in KJ/mol include: Hydrogen bonds, electrostatic energy, Van der Waals energy, and total stabilizing energy. PPCheck, can also predict reliably the correct docking pose by checking if the normalized energy per residue falls within a standard energy range of -2kJ/mol to -6kJ/mol which was obtained by studying a large number of well characterized protein-protein complexes [30]. Additionally, the percentage of residues in the interface of each of the dimers, at the different conformations, was investigated using the PDB-PISA server.

Analysis of the following conformations: 0, 20, 60, 100, 120, 160 and 170 ns, was carried out employing the mentioned servers and are summarized in Tables S1 and S2. A general observation is that the number of interface residues in the different conformations of β_1_-β_1_ dimers vary from that of β_2_-β_2_ dimers. This difference tends to be remarkable in the earlier protein conformations of the molecular dynamics simulations (Table S1).

The majority of the conformations (0, 20, 60, 100, 120 ns) of β_1_ dimers show lower stabilizing energy in comparison to the conformations for β_2_ dimers. The normalized energy per residue is around -2 kJ/mol in various conformers of β_1_-β_1._ While in β_2_-β_2_ none of the conformers reached that value. Thus, suggesting that in general, β_1_-β_1_ dimer is more stable in comparison to β_2_-β_2_ dimer (Table S2). These analyses, comparing selected snapshots of the trajectory do not seem to reflect the dynamic characteristics of each interface. Therefore, we used other tools in order to analyze and compare the dimeric interfaces.

#### 2.4.1 Searching for hot spots within dimeric interfaces

Protein-protein interactions in the interfaces were calculated through PDBsum. Figure 4 depicts protein-protein interactions of the conformers of β_1_-β_1_ and β_2_-β_2_ taken at different times: 0,100,120 and 160 ns. The residues in the interface are depicted and some residues show to be constant in the interface of most of the conformers, for β_1_-β_1_ : Lys173, Gly225, Asn226, Glu228, Thr264, Leu266 and for β_2_-β_2_: Arg130, Thr155, Ile163, Asn220 which are therefore considered the hot spots residues.

**Figure 4.**
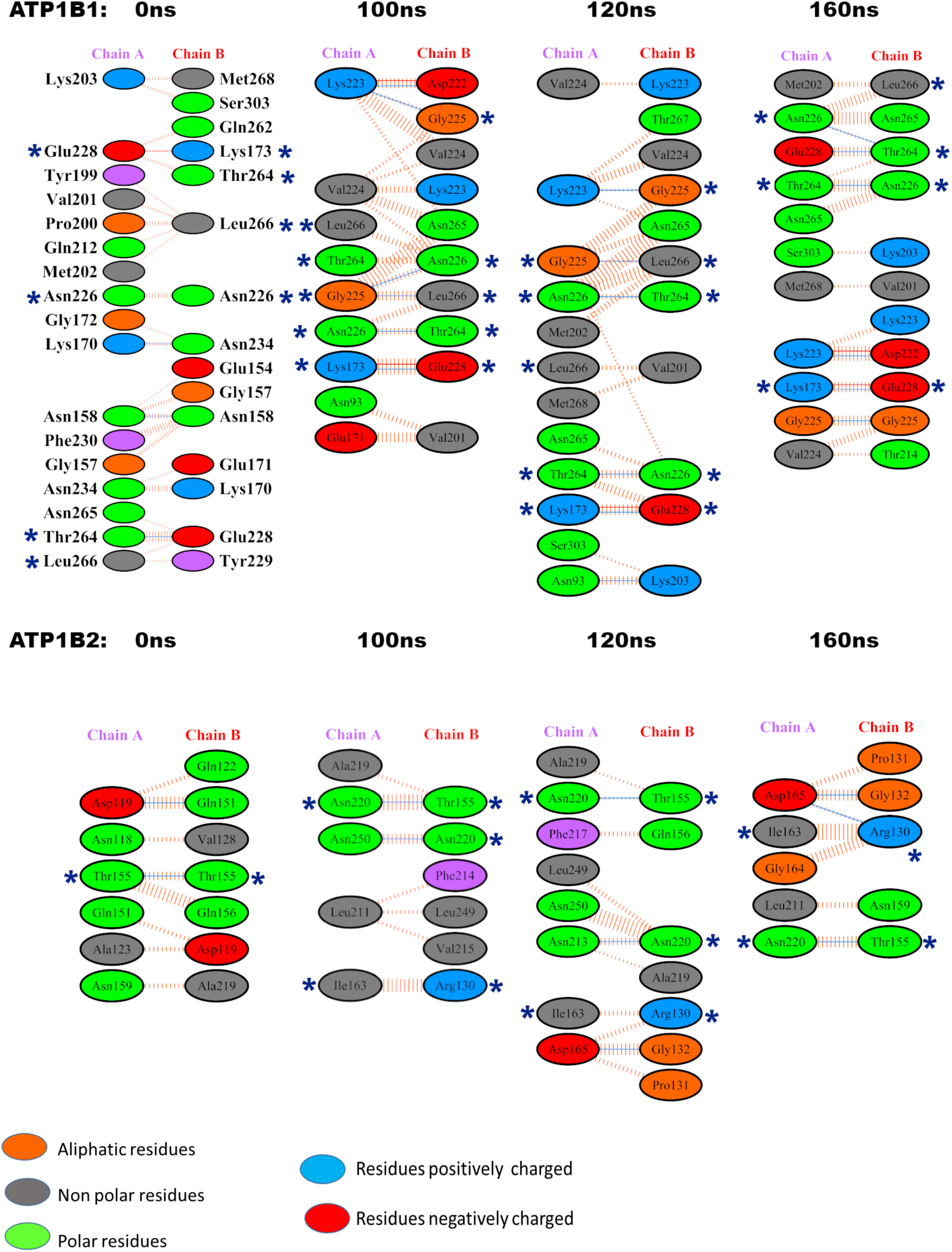
Interfaces at ATP1B1 and ATP1B2 dimers: Graphical representation of the protein interfaces at β_1_ and β_2_ dimers in different snapshots obtained from the MD simulations using PDB SUM server. (*) Residues that appear in all conformations are marked as hot spots.

The analysis of hot spot residues in each dimer suggests significant differences in the interfaces involved in homophilic protein-protein interactions between the β_1_ and β_2_ subunits of Na+,K+-ATPase across neighboring cells. The multiple sequence alignment presented in Figure 5 indicates that, despite high homology between the two subunits, the surface regions engaged in trans-dimerization differ. Notably, the hot spot residues of the β_1_ dimer are clustered in close proximity, while those of the β_2_ dimer are more widely dispersed across the surface. In this alignment, we compared the sequences of the dog ATP1B1 interface, as described in references 15 and 16, with those of human ATP1B1 and ATP1B2 examined in this study. Our findings reveal that: i) β_2_ lacks segment 1 present in both dog and human β_1_ (indicated by the orange box); ii) human β_1_ shares hot spot residues with dog β1 (highlighted in the green box); iii) residues Glu228, Lys173, Thr264, and Leu266 are conserved in both ATP1B1; and iv) residues Gly225 and Asn226 are identical across the three sequences. In relation to the hot spot residues of ATP1B2, we found that i) Arg130, Thr155, and Asn220 exhibit a lack of conservation; ii) although Ile163 is identical across the three sequences, it is exclusive to the β_2_-β_2_ interface. Additionally, while human ATP1B2 shares Gly225 and Asn226 within segment 2, these residues do not seem to influence the β_2_-β_2_ interface. Collectively, these structural distinctions provide insights into the observed differences between the dimer interfaces of β_1_ and β_2_.

**Figure 5.**
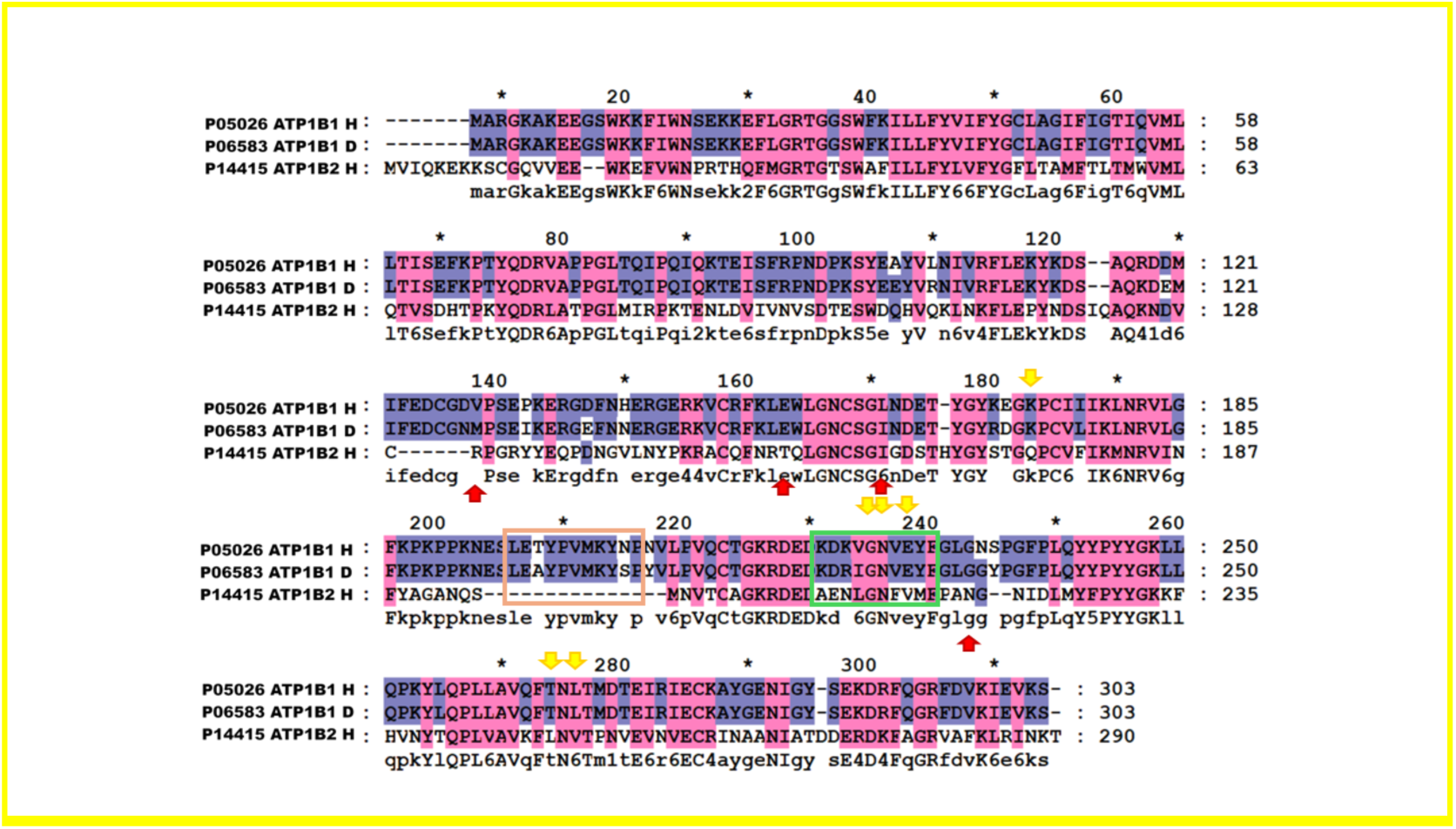
Multiple sequence alignment of ATP1B1 and ATP1B2. A multiple alignment is shown (ATP1B1 Human (P05026), ATP1B1 Dog (P06583), and ATP1B2 Human (P14415). Yellow Arrows: Hot Spots residues of human ATP1B1. Red arrows: Hot spots residues of human ATP1B2. Orange box: Dog Sequence 1 from ref. 15 and Green box: dog Sequence 2 from ref. 16.

##### Monitoring interactions that involve the hot spot residues

Fersht and coworkers provided valuable information regarding the role of hydrogen bonds in protein stabilization [31]. Afterwards, several experimental studies were carried out on proteins of different nature, for example: BPTI [32], RNase Sa [33], Staphylococcal nuclease [34], human lysozyme [35]. In this work, one of the aims was to get insights about hydrogen bonds that are located in β_1_-β_1_ and β_2_-β_2_ dimers. For the case of β_1_, from the three hot spot residues we could identify the formation of hydrogen bonds between the residues: Asn226A and Thr264B (Fig. 6A). On the other hand, the hot spot residues in β_2_-β_2_ identified as forming hydrogen bonds are Asn220A and Thr155B.

**Figure 6.**
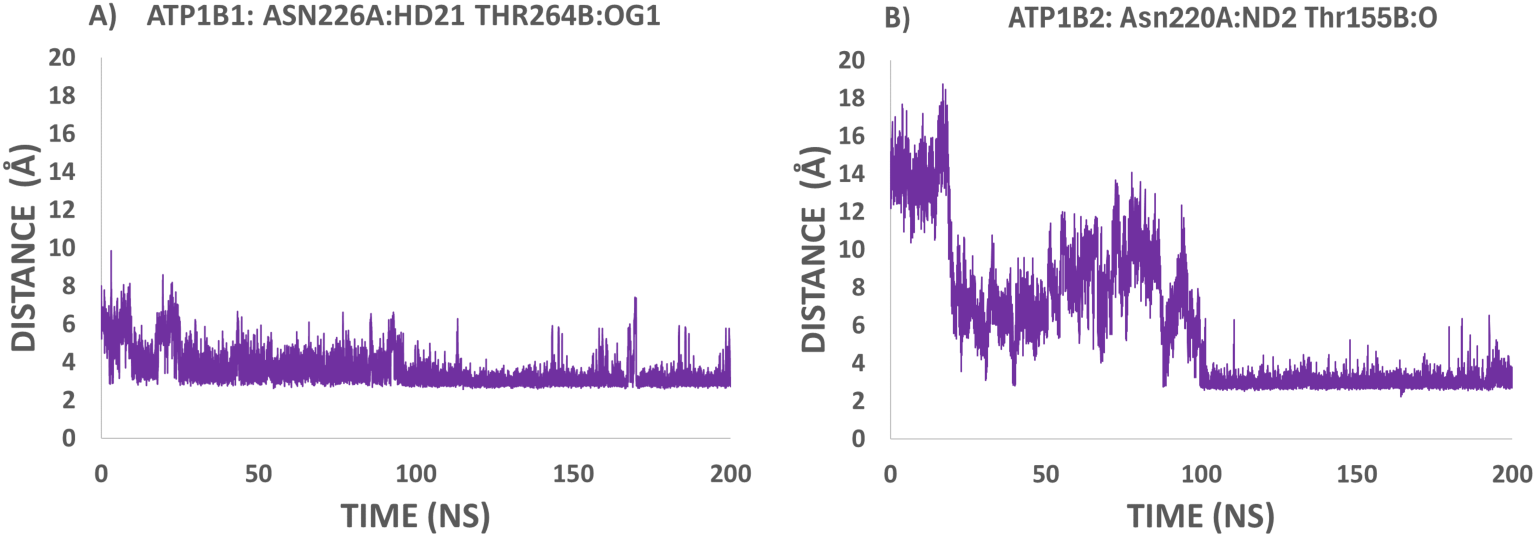
Constant hydrogen-bonds in β_1_ and β_2_ dimers. Distance between atoms (Asn226A:HD21-Thr264B:OG1) for β_1_ and (Asn220A:ND2-Thr155B:O) for β_2_ were calculated along the trajectories using Carma Software.

These Hydrogen bonds were monitored along the trajectories of both β_1_-β_1_ and β_2_-β_2_ dimers. Figure 6A describes a constant hydrogen bond between residues Asn226A and Thr264B in β_1_ dimer, since the first 20 ns of simulation. Figure 6B shows the formation of the hydrogen bond between Asn220A and Thr155B in β_2_ dimer just after 100 ns of simulation. These results indicate a clear difference in the dynamics of the dimers formation and strongly suggest that these hydrogen bonds play a key role in the stabilization of the dimer.

#### 2.4.2 Participation of N-glycosylated residues in the dimeric interface

N-glycosylation involves the attachment of N-acetylglucosamine (GlcNAc) to the nitrogen atom of an asparagine (N) side chain. This type of glycosylation is prevalent in many human proteins and is important for protein folding and stability of the protein [36], and targeting specific cellular locations [37,38]. In the case of β-β interaction it was observed that cell-cell adhesion is impaired when N-glycosylation is inhibited [39,24]. As already known, the main structural difference between β_1_ and β_2_ isoforms is their extent of N-glycosylation sites, β_1_carries three sites while β_2_ carries seven sites.

Here, we identified the N-glycosylation sites located in the interfaces of β_1_-β_1_ and β_2_-β_2_ and their interactions with the surrounding residues at different conformers obtained from the MD simulations (0, 20, 60, 100, 120, 160, 170 ns). Tables 3 and 4 summarizes the interacting glycans of N-glycosylated residues in each dimer and are centered in the table and labeled in red. For the case of the glycosylated Asparagines in β_1_-β_1_ dimer (Table 1), all the identified interactions are intramolecular, with residues of the same chain and are mainly through Van der Waals interactions. The most frequent intramolecular interactions with the glycosylation of Asn265 is through Van der Waals interactions, although few intramolecular hydrogen bonds were identified within the B chain (Thr267B y Thr270B). Noteworthy is the interaction with Thr264 considered a hot spot residue in β_1_-β_1_ interface.

**Table 1.**
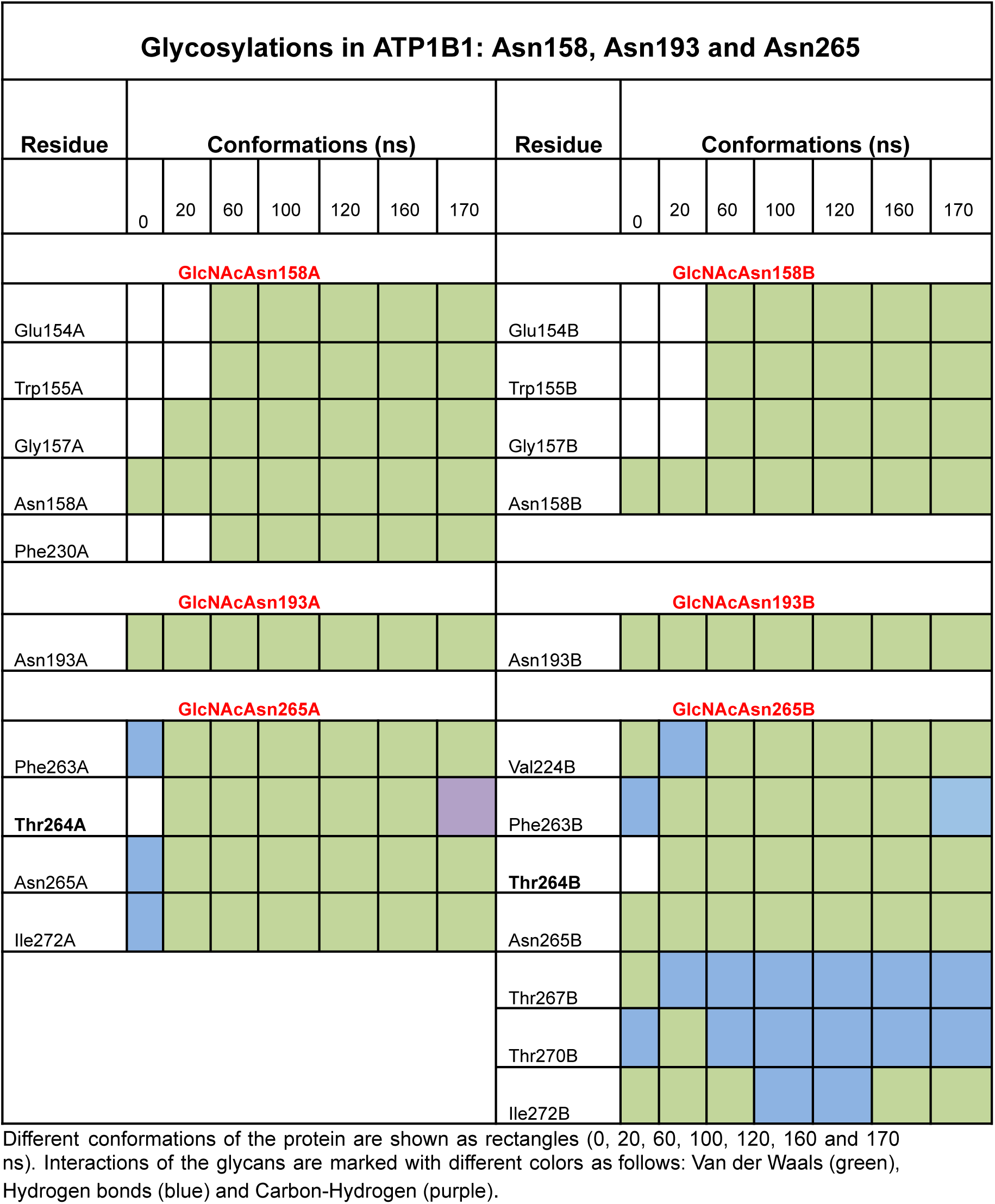
Glycan-protein interactions of ATP1B1 (β_1_ dimer)

Table 2 shows the most frequent interactions with the glycans in the interface of the dimer β_2_-β_2_. Two glycosylated residues, Asn118 and Asn197, show few interactions, mainly intramolecular ones. The other two are more interactive. Asn153A interacts with residues of the same chain mainly through Van der Waals interactions. Nevertheless, Asn153B interacts with residues of the contrary chain through Van der Waals and hydrogen bonds; a similar behavior is observed with Asn159A and Asn159B. Of worthy interest, residues Arg130B and Thr155A which we identified as hot spots within β_2_-β_2_ interface, interact with the glycans of Asn159A and Asn153B, respectively .

**Table 2.**
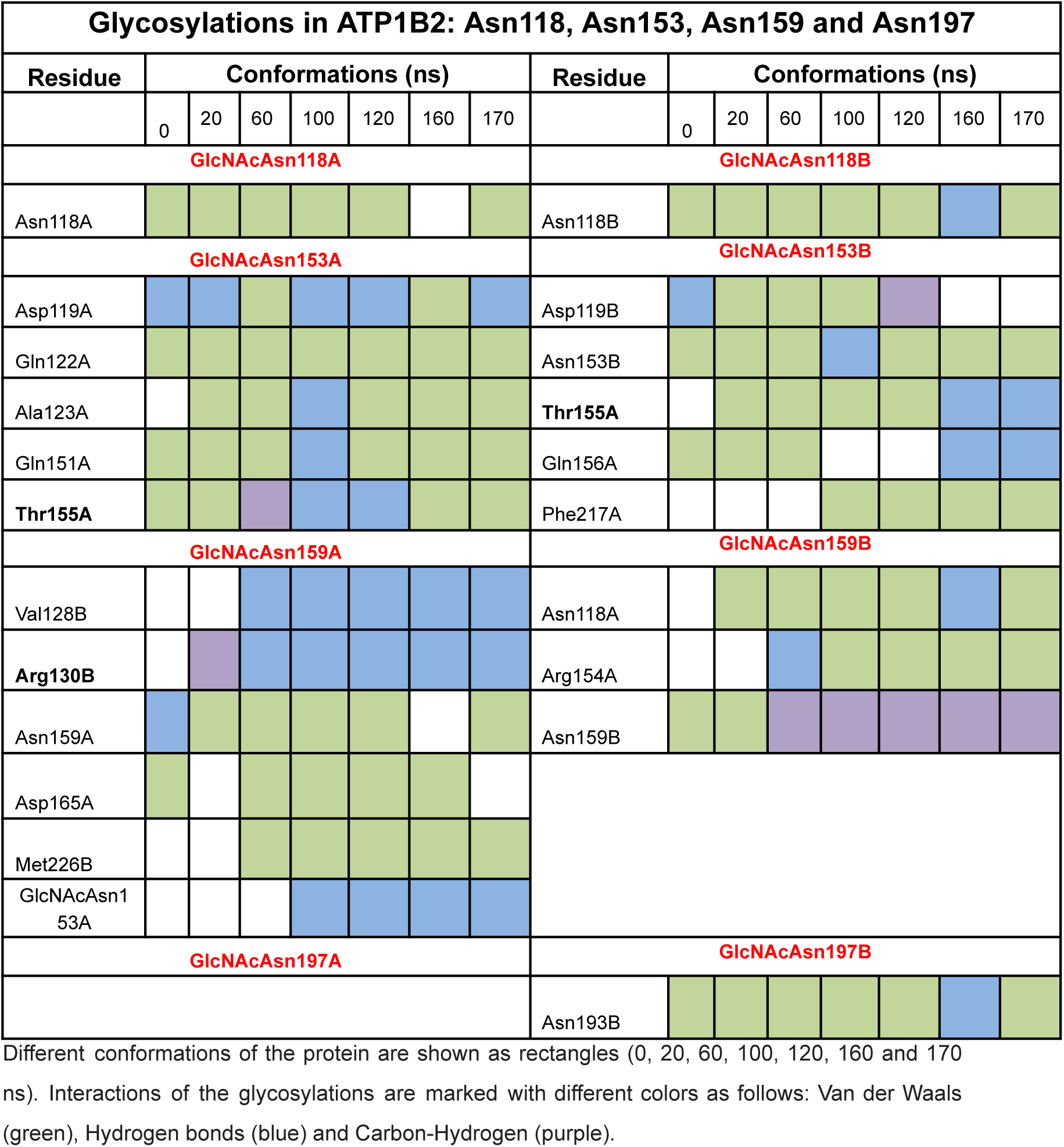
Glycan-protein interactions of ATP1B2 (β_1_ dimer)

**Table 3.** MM-PBSA calculation of ATP1B1 (β_1_) and ATP1B2 (β_2_) dimeric complexes.

| Protein | Electrostatic (kcal/mol) | Van der Waals (kcal/mol) | Poisson-Boltzmann (kcal/mol) | Surface Area (kcal/mol) | Gas (kcal/mol) | Solvate (kcal/mol) | Polar interactions (kcal/mol) | Non Polar interactions (kcal/mol) | Total (kcal/mol) |
| --- | --- | --- | --- | --- | --- | --- | --- | --- | --- |
| ATP1B1 | -11375.42 | -1664.37 | -9782.35 | 151.007 | -13039.79 | -9631.34 | -21157.77 | -1513.361 | -22671.1333 |
| ATP1B2 | -9546.50 | -1514.63 | -8784.48 | 138.04 | -11061.14 | -8646.44 | -18330.98 | -1376.600 | -19707.5810 |

### 2.5 Prediction of Binding Free energy through MM-PBSA method

Here we present an easy-to-use pipeline tool named Calculation of Free Energy (CaFE) to conduct Molecular Mechanics Poisson-Boltzmann Surface Area (MM-PBSA) and LIE calculations. Powered by the VMD and NAMD programs, CaFE is able to handle numerous static coordinate and molecular dynamics trajectory file formats generated by different molecular simulations. The MM-PBSA approach has been widely applied as an efficient and reliable free energy simulation method to model molecular recognition, such as for protein-ligand binding interactions [40]. Moreover, MM-PBSA and MM-GBSA methods are useful methods in both accuracy and computational effort between empirical scoring and strict alchemical perturbation methods [41].

Binding free energy of the dimer complexes β_1_ and β_2_ was calculated by CaFE to conduct MM-PBSA [42] and the obtained values are presented in Table 3. It can be seen that the major contribution to the free energy of the complex is due to polar interactions. Dimeric complex of β_2_ shows higher binding free energy in comparison to the dimeric complex of β_1_ (-19707.5 vs -22671.13 kcal/mol) .

### 2.6 Principal component analysis (PCA)

*In Silico* approaches are useful to describe protein dynamics, in which fluctuations range from bond-distance variations. Molecular dynamics simulations along mathematical applications are very helpful to investigate these fluctuations that occur in the proteins.

Principal component analysis (PCA) is a useful mathematical technique to reduce a multidimensional complex set of variables to a lower dimension. This technique has been used to investigate the stages of protein folding in proteins of diverse nature [43].

In general, the great majority of proteins show particular behavior in which their two/three principal components describe the main motions of the proteins (about 70-80%). As we can infer from our results, the cumulative contribution to the variance in the conformational space is the largest for the first two principal components, 50% and 30% for β_1_ and β_2_ respectively (Figure S1). Dihedral angle principal component analysis (dPCA) has shown advantages for the treatment of proteins and was therefore used for this study .

We studied the fluctuations of these principal components (PC1 and PC2). Projection of the trajectories onto PC1 and PC2, together with the cluster analysis (where the most populated region is highlighted as cluster 1) is depicted in figure S2. We observed that β_1_ homodimer presents low values for PC1 (around 0) and high values for PC2 (around 5) in its main cluster, which is rather localized. On the other hand, the main cluster for β_2_ homodimer spans a large region where values for PC1 range from -6 to -2 and values for PC2 range from -4 to 1. Having a principal component close to zero as is the case for β_1_ dimer suggests the formation of a stable interface that has a short-ranged oscillation. On the other hand, large absolute values in the principal components, as observed for β_2_ dimer, suggest that under these conditions a stable dimer is not yet reached.

The free energy landscapes (Fig. 7) obtained from the dPCA analysis, reveal that for β_1_ dimer the most populated region (ΔG=0) is rather localized, with low values for PC1 (around 0) and high values for PC2 (around 5), whereas for β_2_ dimer the basin where ΔG=0 spans a large region, where values for PC1 range from -6 to -2 and values for PC2 range from -4 to 1. The behavior observed for ATP1B1 suggests that a stable interface is reached where motion tends to stop (PC1=0) and oscillates with a rather small amplitude (-1<PC1<1). In contrast, the large intervals observed for both PC1 and PC2 in ATP1B2 point to the lack of a stable interface.

**Fig. 7.**
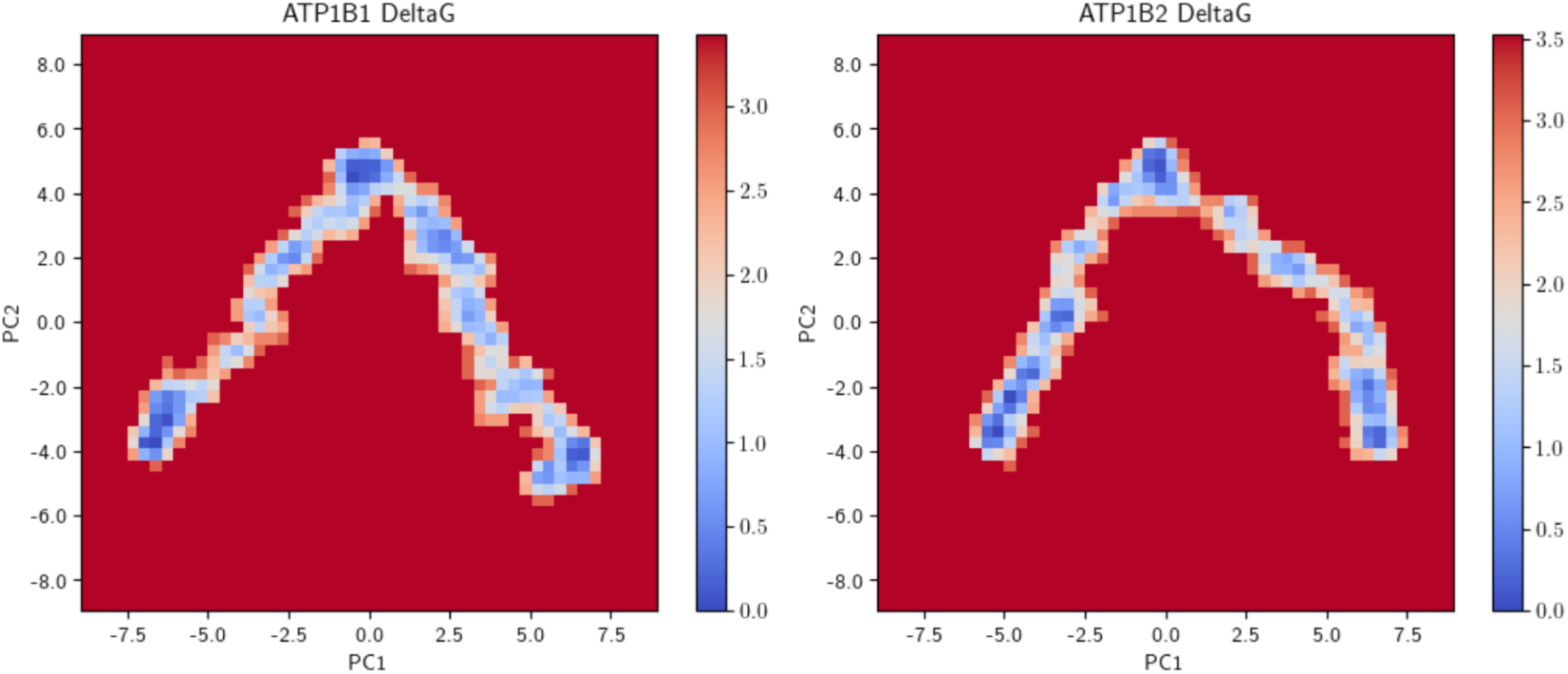
Free energy landscapes considering the first two principal components of β_1_-β_1_ and β_2_-β_2_ dimers.

The motions associated with PC1 and PC2 for β_1_ dimer both show symmetric, rotatory behaviors, whereas, PC1 for β_2_ dimer shows a longitudinal motion and PC2 seems to involve substantially the most mobile loops; this participation of the loops in PC2 suggests that a concerted motion involving the interface has not been reached for this complex (Figure 8 and Supplementary Videos).

**Figure 8.**
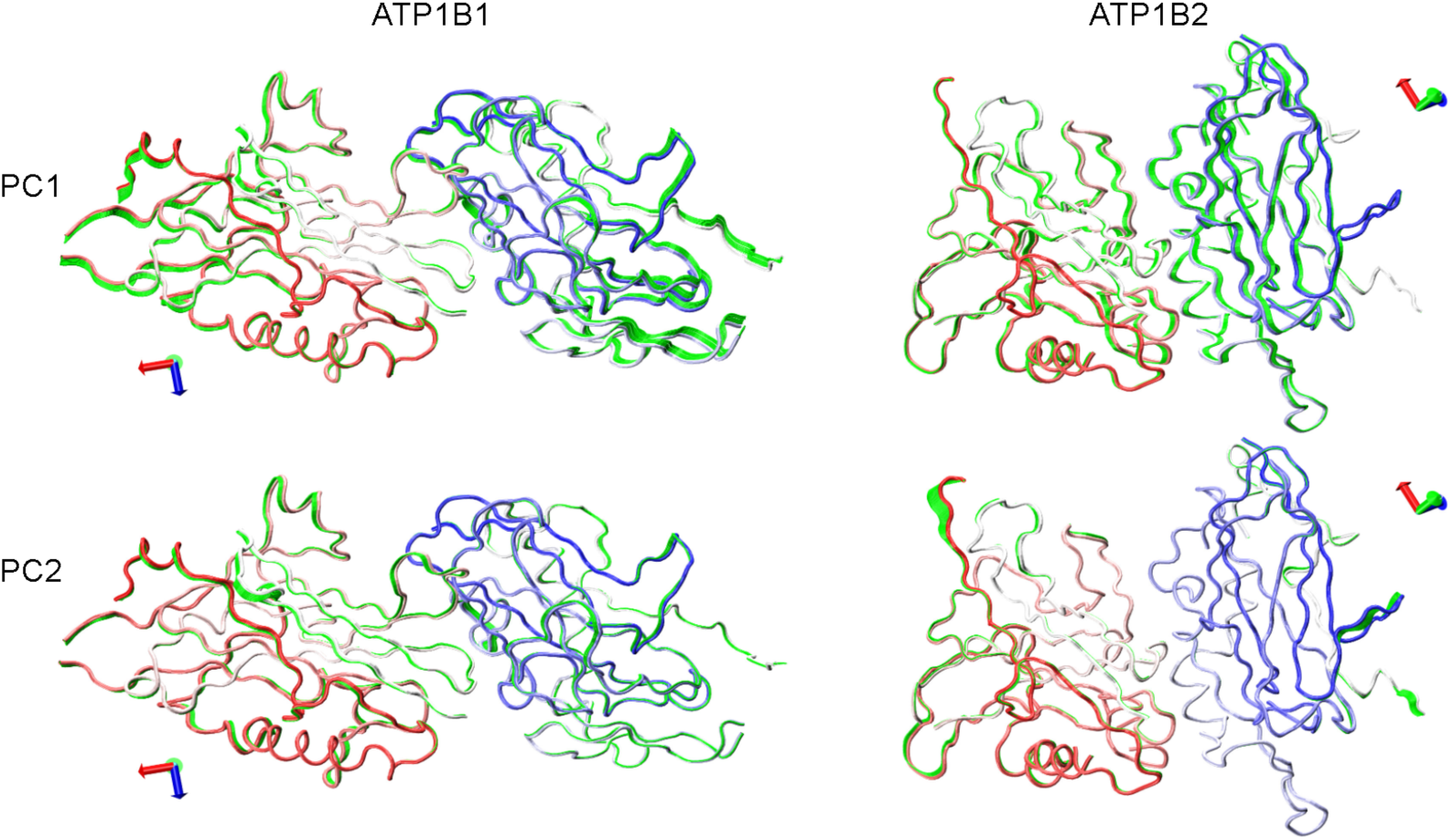
Motion associated with PC1 and PC2. Chain A goes from red in the N-terminus to white in the C-terminus while chain B goes from white in the N-terminus to blue in the C-terminus. Green tubes show the motion associated with the principal components.

### 2.7 Analysis of the movement contributions per dihedral angle

In the dPCA, each dihedral angle *γ* is transformed into a space with two coordinates (cos*γ*, sin*γ*). Each principal component has a weight calculated for each of those coordinates and a measure of the influence of angle *γ* on principal component k (Δ ^(k)^) is defined as the sum of the squares of the corresponding weights, as detailed in section 4.8. The contribution to the first principal component from every angle (Δ^(1)^) was calculated (Figure S3).

For β_1_, elevated contributions are observed in the vicinity of interface residues Val129, Pro130, Glu132 and Pro133 in Chain A, and in the vicinity of interface residues Glu165, Thr166, Asp218 and Asp220 in Chain B. Thus, all interface residues are close to regions with a large contribution and are therefore participating in the main motion of the complex. These observations suggest a stable interface for this complex.

On the other hand, the motion in β_2_ is quite asymmetrical, while there are major peaks for interface residues Asn193 and Ala265 in both Chain A and Chain B, interface residues Met216, Ala219, Asn220, Gly221, Asn222, Ile223, Asp224 and Lys234 are in a region with a rather small contribution in Chain A and regions with small to negligible contributions in Chain B. Interface residue Gly158 is in a region with almost zero contribution in both Chain A and Chain B. The contribution of residue Asn197A in β_2_ to the motion is high, whereas Asn197B does not move significantly, probably due to other interactions that restrain this movement. These findings suggest that the interface is not part of the main motion of this complex and is thus not likely to have reached a stable state.

## 3. DISCUSSION

Our in silico investigations yield significant insights into the structural dynamics underlying the trans-dimerization process of the extracellular domains of human Na^+^,K^+^-ATPase β-subunits, namely *ATP1B1* (β_1_) and *ATP1B2* (β_2_). Previous works have individually studied the structural features of Dog β_1_ [15,16,25] and Human β_2_ [24], revealing notable molecular and biological distinctions in their adhesive properties. In the current study, we expand upon the analysis of β_1_ and β_2_, employing docking and molecular dynamics (MD) simulations and leveraging *in silico* methodologies to investigate various structural aspects. Our aim is to elucidate the biological disparities observed between β_1_ and β_2_/AMOG subunits as adhesion molecules.

### Analysis of the interacting interfaces of β_1_-β_1_ and β_2_-β_2_ dimers along the MD trajectories

Initially, we examined the interacting interfaces of both β_1_-β_1_ and β_2_-β_2_ dimers, revealing a consistent reduction in the number of residues within the interface of *β_2_-β_2_* dimer throughout the simulation compared to β_1_-β_1_ dimer (see Fig. 4). This reduction in residue count corresponds to a lower interface area in β_2_-β_2_ dimer, which correlates with reduced complex stability attributable to weaker interactions within this region (Table 5). Furthermore, we identified hot spot residues pivotal for interface stabilization. Notably, the number of hot spot residues within the β_2_-β_2_ interface was found to be lower than that within the β_1_-β_1_ interface.

The interface of the dimer of human β_1_-β_1_that represents the starting point for the molecular dynamic simulation (0 ns in Figure. 4) is very similar to that of the dog β_1_-β_1_ proposed by the *in silico* and *in vitro* analyses of [16]. Nevertheless, during the MD simulation the interactions at the interface become less ample and at least 5 residues are identified as hot spots localized in a domain that range between Gly225, and Leu266. On the other hand, the hot spots residues of β_2_-β_2_ dimer are more dispersed and indicate a large interface surface. The alignment of *human and dog ATP1B1* and human *ATP1B2* in figure 5 shows that indeed the apparent hot spot residues are localized in very distant domains on β_2_-β_2_ dimer that do not overlap with those of β_1_-β_1_ interface. This observation correlates with the MM-PBSA calculations (Table 5), that shows lower binding free energy for β_2_-β_2_ dimer in comparison to β_1_-β_1_ (-19707.5 vs -22671.13kcal/mol, respectively).

Meticulous analyses of the first two principal components from the dPCA performed revealed significant differences in the dynamics of the β_1_ and β_2_ dimers. First, the most populated energy region for β_1_ is rather localized whereas for β_2_ the most populated energy region spans a large area. Besides, β_1_ shows a symmetric, concerted rotation involving the interface residues in both monomers, whereas β_2_ shows a tendency to increase the distance between the monomers in its main motion (PC1), while the second main motion does not involve significantly the interface residues. Additionally, for β_1_, an important contribution is observed for the dihedrals surrounding Asn158 and Asn193. The former is close to interface residues Glu165 and Thr166 and its pronounced motion could influence this region of the interface. Interface residues Asp218 and Asp220 show important contributions, which suggests their involvement in the concerted motion of the interface. In contrast, most glycosylated asparagines and interface residues show minute contributions for β_2_ dimer. All these observations are consistent with the existence of a stable interface in β_1_ dimer and the lack thereof in β_2_ dimer. Interestingly, for β_2_ dimer Asn159A, Asn153B and Asn159B form favorable interactions with residues in the opposite chain (Table 4), which suggests that these glycosylations could play an instrumental role in keeping the dimer despite the motions that tend to separate the monomers.

### Biological divergence observed between β_1_ and β_2_/AMOG subunits

Drawing upon structural analysis, stability assessments, movement analyses, and free energy calculations conducted in this investigation, it is elucidated that β_2_, a conventional constituent of astrocytes Na^+^-pump, does not engage in β_2_-β_2_ *trans*-dimerization among astrocytes [17], neither in vitro as evidenced by protein-protein interaction assays such as pull-down. However, a contrary trend is observed when β_2_ is expressed in CHO and MDCK cells [24]. These findings imply the presence of additional modulatory elements that either facilitate (in CHO or MDCK cells) or hinder (in astrocytes) the formation of stable β_2_-β_2_ *trans* interactions. Notably, the potential *cis* interactions of surface residues of the protein with those from the same membrane remain unexplored. These interactions may be masking the *trans* interaction capacity with the β_2_ subunit of adjacent cells.

Moreover, it is pertinent to recall that the Na^+^,K^+^-ATPase, comprising α_2_ and β_2_ subunits, has previously been identified as a constituent of a complex on the astrocytic plasma membrane [44]. This complex has been shown to orchestrate the functional interplay among GluR2, PrP, α_2_-subunit, β_2_-subunit, basigin, and MCT1, thereby tightly regulating lactate transport in astrocytes. In this complex, the N-glycans of β_2_ interact with the lectin domain of basigin [45].

Of worthy interest is that glycan-protein interactions can be considered as multivalent interactions which are often required to achieve biologically relevant binding even though they are known to have low affinity [46]. On the other hand, these interactions have been related to some other functions which include: dynamic forms of adhesion mechanisms, for example, rolling (cells), stick and roll (bacteria) or surfacing (viruses) [46]. Glycosylations play a pivotal role in cell adhesion and recognition, and can also influence protein-protein interactions. Interestingly, the main difference between the β_1_ and the β_2_ isoforms is in their number and sites of N-glycosylation. While human β_1_ (*ATP1B1*) carries three conserved N-Glycosylation sites, β_2_ (*ATP1B2*) conserves these three sites but has 4 additional ones. In the case of cell adhesion mediated by trans-interactions of β_1_-β_1_ and β_2_,-β_2_, the N-glycosylation of both β-subunits had been reported to play an important role [15,24]. Here, in a detailed analysis of the interactions of the glycosylated residues, we observed that this type of interactions in β_1_-β_1_ dimer occur within residues located at the same chain; whereas β_2_-β_2_ dimer shows interactions that occur both intra- and inter-molecular, between contrary chains.

Understanding the cellular physiology of β_2_/AMOG is gaining renewed interest due to the increasing evidence implicating Na^+^,K^+^-ATPase in neurological pathologies and disorders. Aberrant expressions of different Na^+^,K^+^-ATPase subunits and their activity have been linked to the development and progression of various cancers, as well as cancer cell proliferation, migration, and apoptosis [47]. However, the exact mechanism by which Na^+^,K^+^-ATPase influences cellular migration and invasion in cancer remains unclear. In the brain, several mutations and aberrant expressions of Na^+^,K^+^-ATPase α and β isoforms have been associated with both neurological phenotypes [48] and brain cancer [[49]. Remarkably, the majority of Glioblastoma multiforme (GBM) tumors exhibit a dramatic loss of β_2_/AMOG expression. Sun et al. [49] proposed that this loss may be a key mechanism contributing to the increased invasiveness of GBM cells. They found that overexpression of β_2_/AMOG reduced the invasion of GBM cells and brain tumor-initiating cells (BTICs) without affecting their migration or proliferation. Conversely, knockdown of β_2_/AMOG expression in normal human astrocytes increased their invasiveness. Collectively, these findings implicate β_2_/AMOG in glioma invasion, suggesting that downregulation of β_2_/AMOG expression is a crucial step in the differentiation of BTICs. Therefore, β_2_/AMOG is considered a tumor-suppressing protein and is of great interest for understanding its function in the central nervous system.

### Concluding remarks

In this study, we identify structural components that account for some of the differences in the homotypic adhesive functions of β_1_ and β_2_. Firstly, the interface composition varies due to sequence and structural differences. Secondly, the surface glycans differ; there are more N-glycans in β_2_, and while they do not participate in protein-protein interactions in β_1_, they seem crucial for facilitating these interactions in β_2_. Nonetheless, the dimer formed by *trans*-interaction between β_2_ subunits, is not a stable complex, suggesting that a stable β_2_-β_2_ interface requires the participation of additional cellular component(s) (co-factors) that are not taken in consideration in our model. On the other hand, β_2_ subunit is probably able to form a stable interface with a yet unknown neural receptor involved in regulation of neurite outgrowth and cell migration during development. Our future work is directed to identify that heterotypic partner of β_2_ on neurons and study their interaction.

## 4 MATERIALS AND METHODS

### 4.1 Molecular modeling of the monomers of Na^+^, K^+^-ATPase β subunits in humans

Three dimensional (3D) structures of the β-subunits of Na^+^,K^+^-ATPase: *ATP1B1* and *ATP1B2* were obtained by employing the Swiss Model Program [50]. For the *In Silico* studies of both proteins we considered only the extracellular domain of the Na^+^,K^+^-ATPase β_1_ and β_2_ subunit.

The glycosylation and disulphide bridges sites on both *ATP1B1* and *ATP1B2* were taken from the Uniprot database *ATP1B1* (P05026) and ATP1B2 *(P14415)* [52]. For the first protein (β_1_), the following glycosylation sites were considered: N158, N193 and N265, and the disulphide bridges: S126-S149, S159-S175 and S213-S276. For the second protein (β_2_), the following glycosylations were considered: N96, N118, N153, N159, N193, N197 and N238, whereas the following disulphide bridges were considered: S129-S150, S160-S177 and S200-S261. Glycosylations and disulphide bridges of each of the proteins were included by means of the CHARMM-GUI Program [53].

### 4.2 Building the dimers of ATP1B1 and ATP1B2 and validation of the 3D models

Once the monomers were correctly built, the molecular docking of both β_1_-β_1_ and β_2_-β_2_ subunits was performed by using HDOCK server in order to obtain dimer complexes of each of the proteins. HDOCK predicts the interaction of protein-ligand complexes through hybrid algorithm strategy of template-based and template-free docking [53].

After performing protein-protein docking procedure, dimeric complexes were selected according to the criteria a) most energetically favorable β_1_-β_1_ (Docking score -193.04) and β_2_-β_2_ (Docking score -274.99), by means of the HDOCK Server [53], b) trans orientation in the dimeric complexes.

### 4.3 Molecular Dynamics Simulations of dimeric complexes of **β_1_**-**β_1_** and β_2_ -β_2_

MD simulations of both dimers were carried out using CHARMM-GUI Server and considering the commands from the Solution Builder implemented in the mentioned Program. Dimers were in a rectangular waterbox size of 10Å edge distance. A NaCl solution (0.15M) was integrated in the system by using “Distance” as Ion Placing Method [54]. Periodic Boundary Conditions were implemented as Generating grid information for PME FFT automatically. Equilibration of the systems was done using an NVT ensemble and dynamics input was generated as an NPT ensemble (310 K). MD simulations were run for about 200ns.

### 4.4 Differences of the interfaces of the dimeric complexes of **β_1_**-**β_1_** and β_2_ -β_2_

Molecular interactions were analyzed in the different protein conformations which include: hydrogen bonds (kJ/MOL), electrostatic energy (kJ/MOL), Van der Waals (kJ/MOL), and Total stabilizing energy (kJ/MOL). All these parameters were calculated through PPCHECK Software [30] which is a specialized web server useful to identify non-covalent interactions at the interface of protein-protein complexes. Moreover, the percentage of residues in the interface, for both chains (chain A and chain B) was calculated using the Program PDBePISA from the Protein Data Bank in Europe [55].

For this analysis we compared three different conformations at 0ns, 20ns, 60ns, 100ns, 120ns and 160ns.

Protein-protein interactions in the interfaces were calculated through PDBsum (http://www.ebi.ac.uk/thornton-srv/databases/pdbsum/) in which interface areas are computed using Program called NACCESS http://wolf.bms.umist.ac.uk/naccess, which is implemented in the Software.

### 4.5 Prediction of Binding Free energy through Molecular Mechanics Poisson–Boltzmann Surface Area (MM-PBSA method)

Binding free energies of the dimeric complexes of ATP1B1 and ATP1B2 were calculated by means of the pipeline tool named Calculation of Free Energy (CaFE) which is a useful tool to predict binding affinity of some complexes by using end-point free energy methods [42] with the aim to conduct MM-PBSA calculation [40]. In the MM-PBSA analysis, three main energetic components are calculated. Firstly, the gas-phase energy difference between the complex and the receptor separated. Afterwards, the difference of solvent-accessible surface area (SASA) is measured and the non-polar solvation free energy is calculated. Finally the binding free energy is added throughout an ensemble conformations. By means of this analysis we were able to get an insight about non-bonded interactions such as Van der Waals, electrostatic, among other parameters.

### 4.6 Principal component analysis (PCA)

PCA has become a popular method to reduce the dimensionality of a complex system and has been previously applied to G protein-coupled receptors (GPCRs) [56]. This method diagonalizes the two-point covariance matrix, thus removing the instantaneous linear correlations among the atomic positions. It has been shown that a large part of the system’s fluctuations can be described in terms of only a few of the eigenvectors obtained, usually those corresponding to the largest eigenvalues. The principal components are the product of these eigenvectors with the mass weighted coordinates of the molecule and can be used as reaction coordinates and to obtain free energy surfaces of the system, among other analysis that can be performed of this representation of the conformational behavior. We used the dihedral angle principal component analysis (dPCA) version, as modifications in dihedral angles lead often to more dramatic conformational changes than movements in atomic cartesian coordinates [57]. The calculations were performed using the Carma program [58].

### 4.7 Free energy landscapes

The calculated dPCA were used to represent the free energy surface of the system, restricting the surface to two dimensions (thus using the first two principal components V_1_ and V_2_):

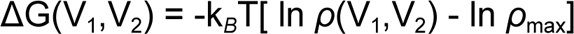

where *ρ* is an estimate of the probability density function obtained from a histogram of the data. *ρ*_max_ denotes the maximum of the density, therefore ΔG=0 for the most populated region [57].

### 4.8 Contribution of movements in each of the residues along the trajectories

In the dPCA, each principal component V_k_ is given by

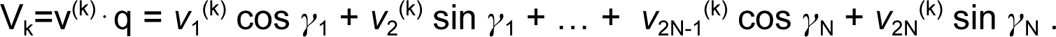

Where v^(k)^ is the k_th_ eigenvector and {*γ*_n_ } n=1,…,N, is the sequence of dihedral angles (*Φ*,*Ψ*) of the peptide backbone.

A measure of the influence of angle *γ*_n_ on the principal component V_k_ may be defined as

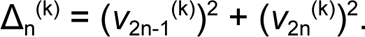

The length of each eigenvector is 1, and thus 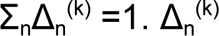 can hence be considered as the percentage of the effect of the angle *γ*_n_ on the principal component V_k_ [57]. These contributions per dihedral angle were calculated for the first principal component 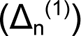 for both ATP1B1 and ATP1B2.

## ACKNOWLEDGMENTS

MMA thanks CONAHCYT for the Ciencia Frontera Project CF-2023-G-1454.

MMA and GRS thank Computing Cluster Xiuhcoatl-Cinvestav (granted by LANCAD) and Hector Manuel Oliver Hernandez.

JLRT thanks SIP-IPN 20240801, SIBE-IPN and EDI-IPN.

## SUPPLEMENTARY MATERIALS

**Table S1.** Interactions in the interface calculated in the different conformations of ATP1B1 and ATP1B2.

| PDB Pisa structure | Percentage of Interface residues |  |  |  |  |  |  |
| --- | --- | --- | --- | --- | --- | --- | --- |
| Protein | Conformation Obtained from MD simulation |  |  |  |  |  |  |
| ATP1B1 | 0ns | 20ns | 60ns | 100 ns | 120ns | 160 ns | 170 ns |
| Chain A | 13.3% | 5.4 % | 8.8% | 5.8% | 7.9% | 7.9% | 7.5% |
| Chain B | 12.1% | 5.8% | 8.3% | 5.8% | 6.7% | 5.4% | 5.4% |
| ATP1B2 | 0ns | 20ns | 60ns | 100 ns | 120ns | 160 ns | 170 ns |
| Chain A | 10.6% | 5.5 % | 6.9% | 11.5% | 8.8% | 9.2% | 7.4% |
| Chain B | 10.4% | 6.5% | 6.9% | 9.7% | 9.2% | 10.1% | 6.0% |

**Table S2.** Interactions* in the interface calculated in the different conformations of ATP1B1 and ATP1B2.

| PPCHECK | Energy of the Interactions in the interface |  |  |  |  |  |  |
| --- | --- | --- | --- | --- | --- | --- | --- |
| **ATP1B1** | **0ns** | **20ns** | **60ns** | **100 ns** | **120ns** | **160ns** | **170 ns** |
| Hydrogen bonds (kJ/mol) | -9.81 | -7.77 | -10.04 | -19.22 | -17.60 | -13.59 | -16.8 |
| Electrostatic energy (kJ/mol) | -12.49 | -14.31 | - 31.94 | -36.42 | - 19.32 | -11.88 | -12.37 |
| Van der Waals (kJ/mol) | -87.19 | -28.09 | -63.6 | -60.33 | -70.07 | -55.09 | -94.19 |
| Total stabilizing energy (kJ/mol) | -109.5 | -50.18 | -105.57 | -115. 96 | -106.98 | -80.56 | -123.42 |
| Normalized energy per residue (kJ/mol) | 1.08 | -0.78 | -1.43 | -2.03 | -1.67 | -1.37 | -2.52 |
| **ATP1B2** | **0ns** | **20ns** | **60ns** | **100 ns** | **120ns** | **160ns** | **170 ns** |
| Hydrogen bonds (kJ/mol) | -10.91 | -14.02 | -9.7 | -11.95 | 0.0 | -11.70 | 0.00 |
| Electrostatic energy (kJ/mol) | 66.98 | 12.36 | - 62 | 3.08 | -43.81 | -34.87 | -4.09 |
| Van der Waals (kJ/mol) | -25.22 | -29.34 | -21.85 | -33.18 | -52.79 | -92.87 | -7.89 |
| Total stabilizing energy (kJ/mol) | 30.84 | -31 | -64.84 | -42.05 | -96.60 | -139.43 | -11.98 |
| Normalized energy per residue (kJ/mol) | 0.33 | -0.72 | -1.10 | -0.55 | -1.40 | -1.81 | -1.11 |

\*The interaction is expressed as pseudoenergy, whose ranges have been standardized using known sets of protein-protein complexes.

**Figure S1.**
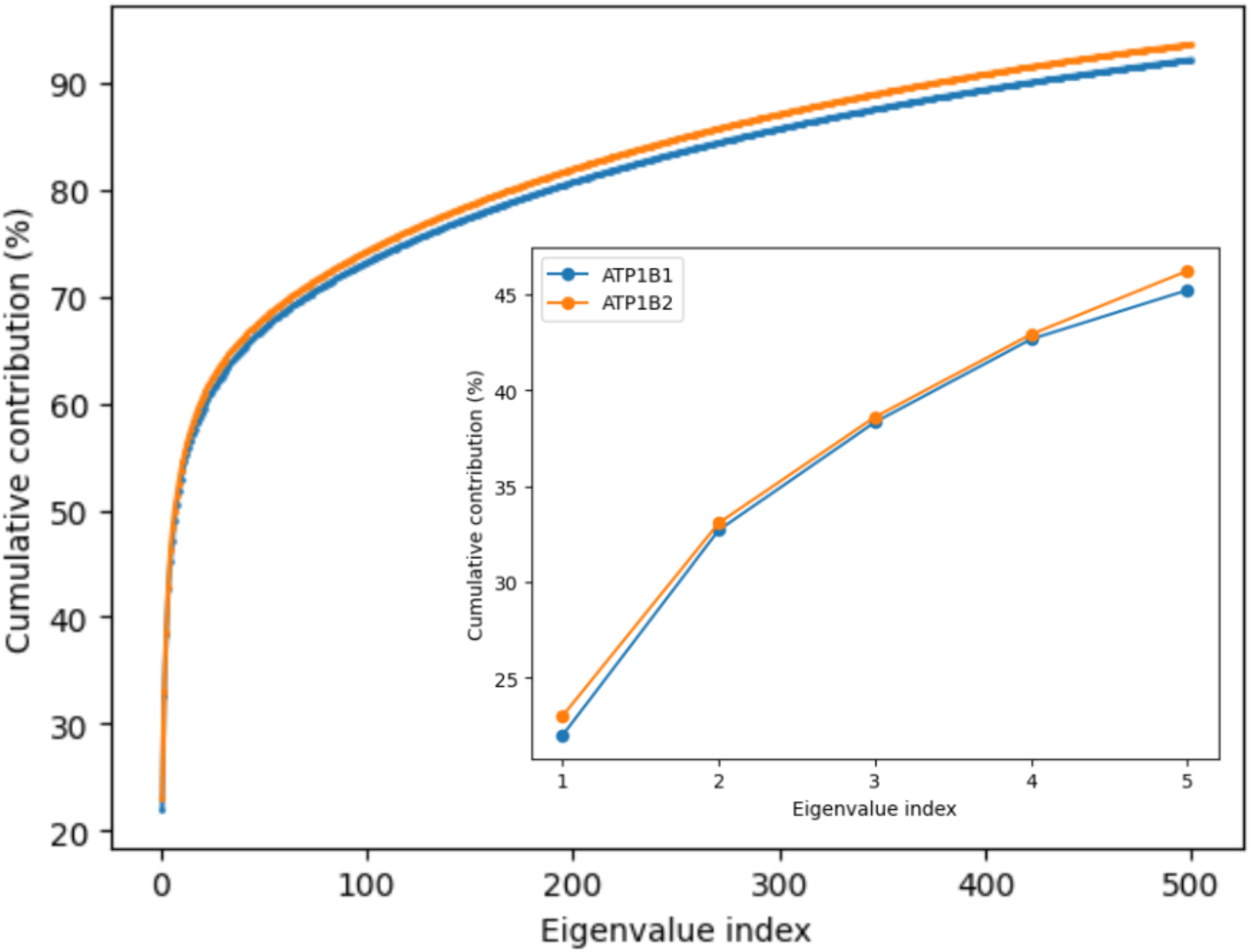
Cumulative contribution of the principal components to the variance in the structural space. The first principal components for both dimers explain under 40% of the energy observed in the simulation. Reaching 80% requires over 100 principal components, which suggests the simulation time might have to be extended to allow fewer motions to dominate the dynamics. The projections of the trajectories onto PC1 and PC2 are very different, which was expected as the sequences show only partial similarity. Regarding the cluster analysis, some clusters of similar size to the main cluster are observed, which correlates with the poor dominance shown by the main principal components and hints at the possibility of a main energetic basin still waiting to be populated. The motions associated with the two main principal components for ATP1B1 show symmetric, rotatory behaviors that are expected in a stable dimer. In contrast, for ATP1B2, PC1 shows an asymmetric, longitudinal motion that seems to drive the monomers away from each other while PC2 seems to involve mostly inconsequential motions in the most mobile loops. This behavior can be related to Table I, where ATP1B2 shows unfavorable interactions for several of the conformations considered

**Figure S2.**
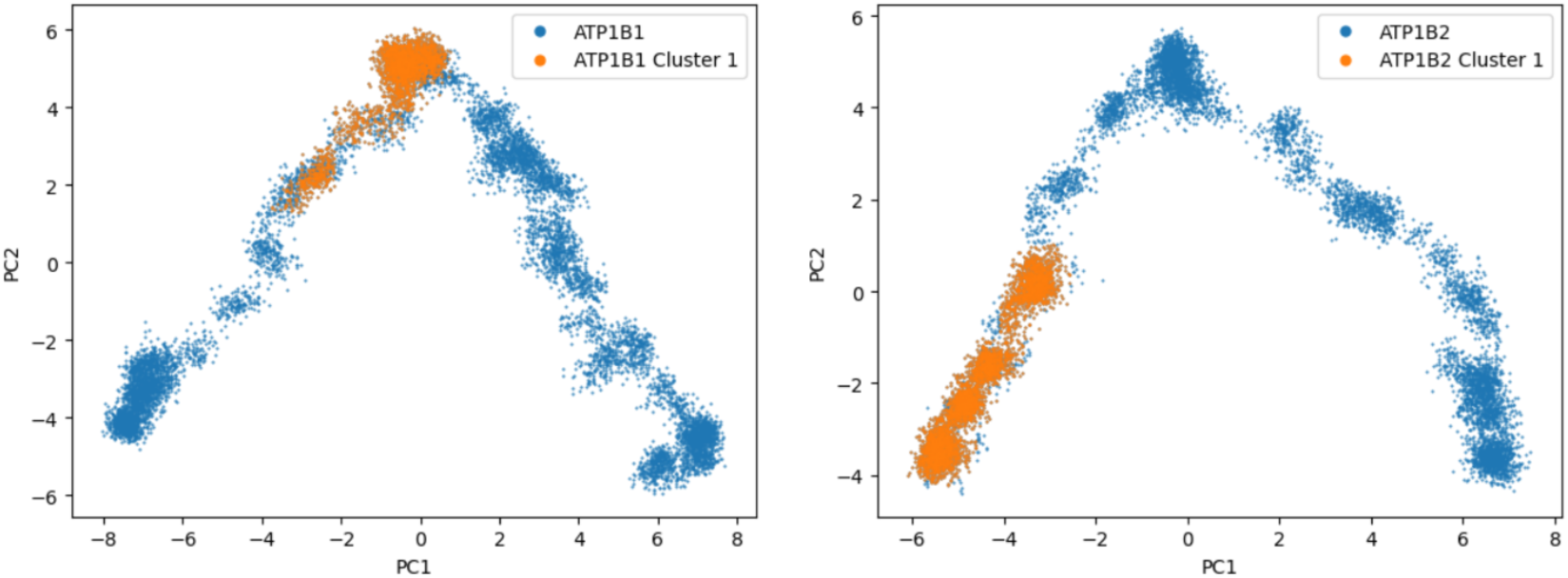
dPCA Cluster analysis considering the first two principal components of β_1_-β_1_ and β_2_-β_2_ dimers. For ATP1B1 we have the main cluster around low values of PC1, whereas the main cluster for ATP1B2 is in a region with large values for both PC1 and PC2, and it also spans a larger region. These observations support the conclusion that ATP1B1 shows a stable interface and the lack thereof for ATP1B2.

**Figure S3.**
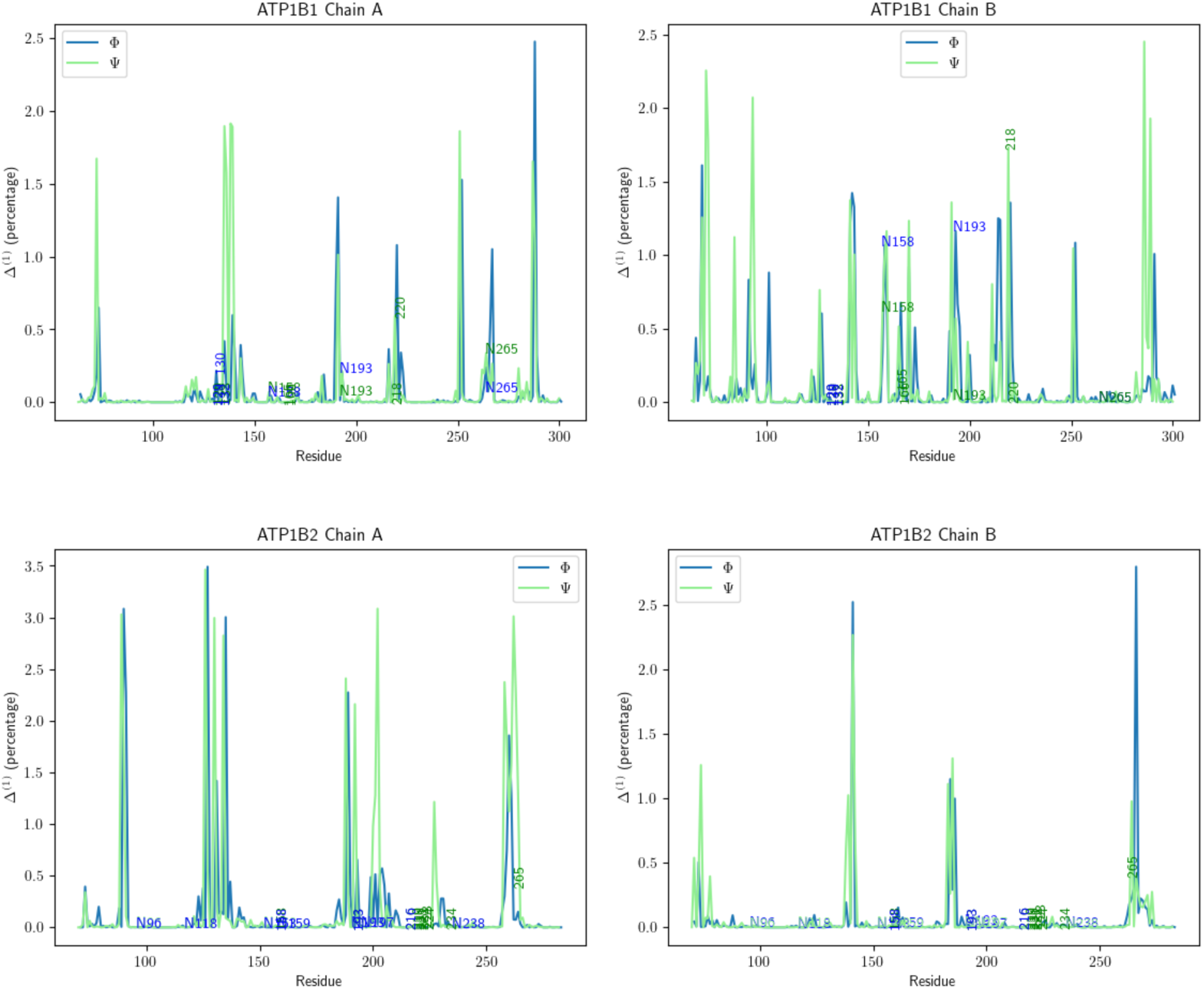
Analysis of the movement contributions per dihedral angle. Vertical numbers indicate interface residues while horizontal numbers indicate glycosylated asparagines. For ATP1B1, Chain A, the highest peaks are in 5 𝚽 angles (Asn193, Asp222, Tyr254, Asp269, Arg290) and 9 𝚿 angles (Val72, Glu135, Arg136, Asp138, Phe139, Asn193, Lys253, Asp289, Arg290), while for Chain B the highest values are found in 10 𝚽 angles (Tyr68, Arg143, Gly144, Glu145, Ser160, Ser195, Lys216, Arg217, Asp222, Tyr254) and 15 𝚿 angles (Tyr68, Asp70, Arg71, Gln84, Asn93, Arg143, Glu145, Gly161, Gly172, Asn193, Lys221, Lys253, Lys288, Phe291, Gly293). The distribution of peaks for ATP1B2 is somewhat different, as for Chain A the highest peaks are in 8 𝚽 angles (Asn90, Leu91, Cys129, Arg133, Gln137, Asn193, Ala266, Asn267) and 17 𝚿 angles (Glu89, Asn90, Val128, Gly132, Glu136, Ala192, Met196, Asp205, Glu206, Tyr231, Asn264-Thr270) while for Chain B the highest values are in 3 𝚽 angles (Leu143, Phe188,Asp272) and 5 𝚿 angles (Gln74, Gly141, Leu143, Asn187, Tyr189).

**Video S1 dPCA_5.**
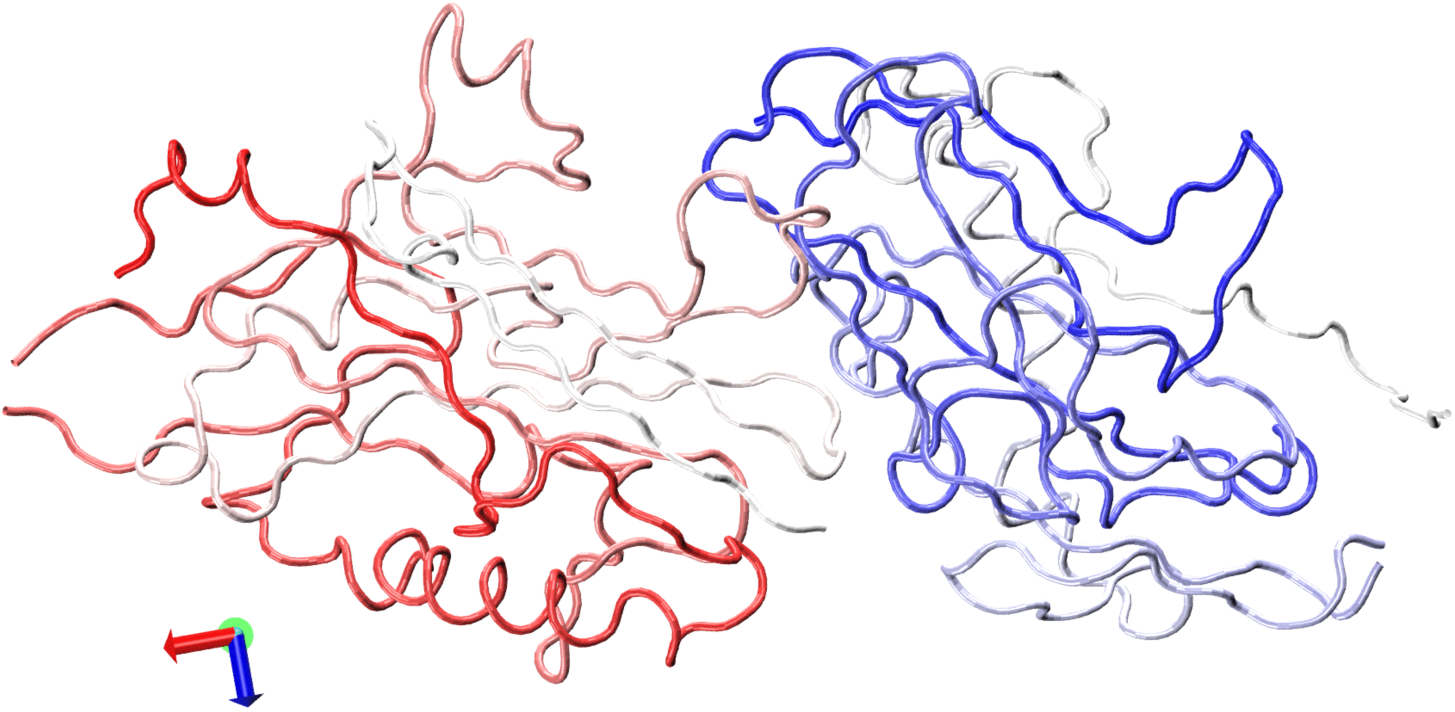
Motion associated with ATP1B1 dimer, PC1.

**Video S2 dPCA_5.**
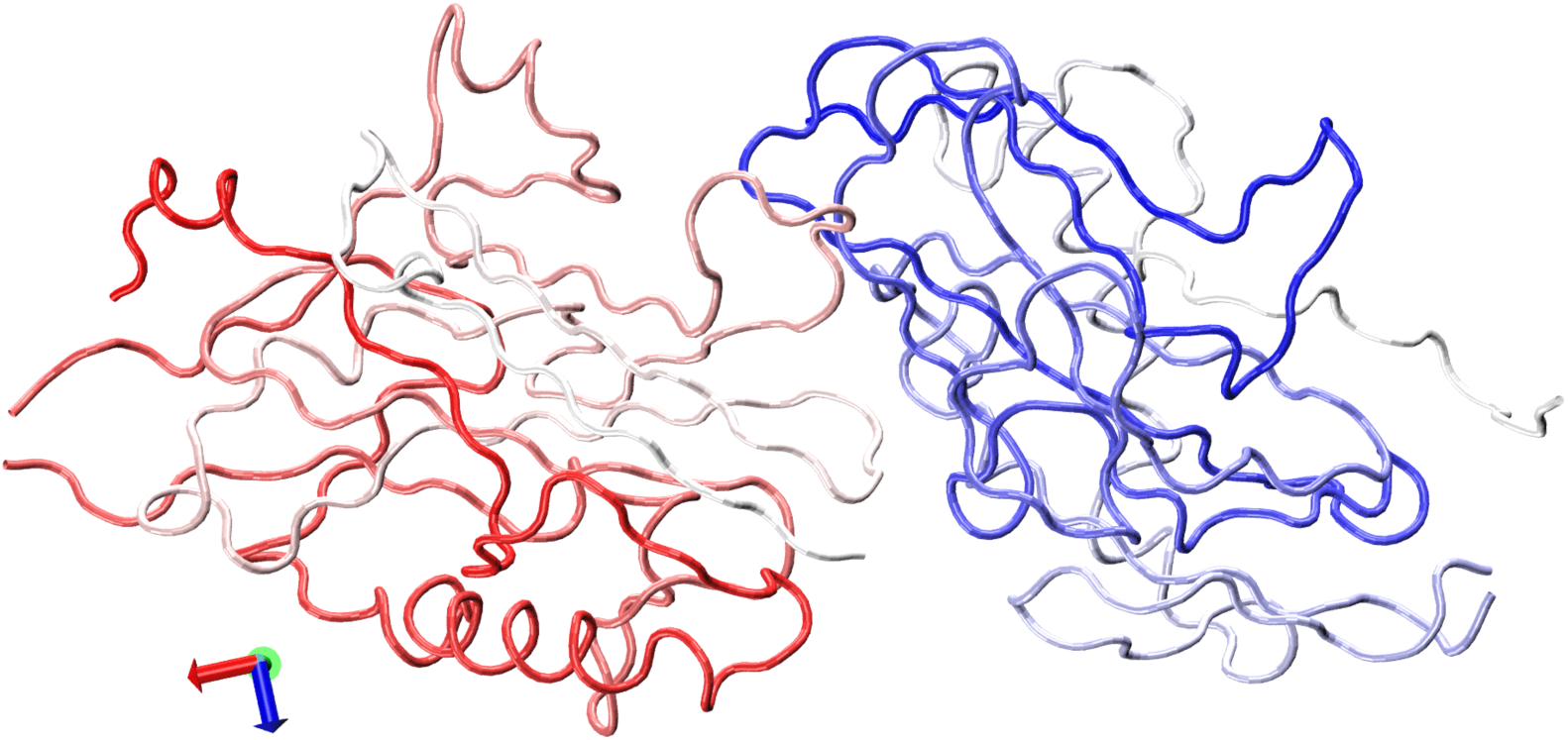
Motion associated with ATP1B1 dimer, PC2.

**Video S3 dPCA_5.**
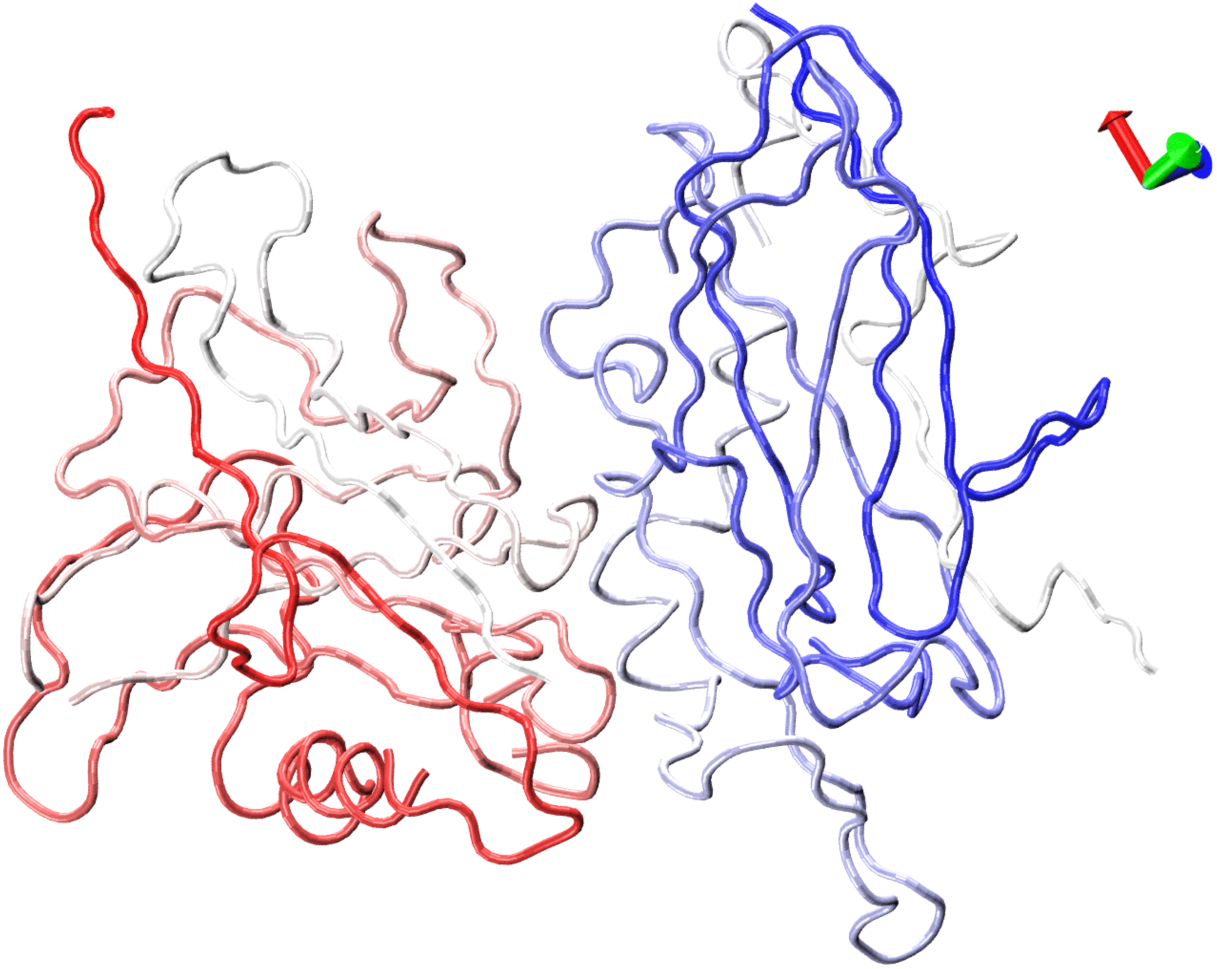
Motion associated with ATP1B2 dimer, PC1.

**Video S4 dPCA_5.**
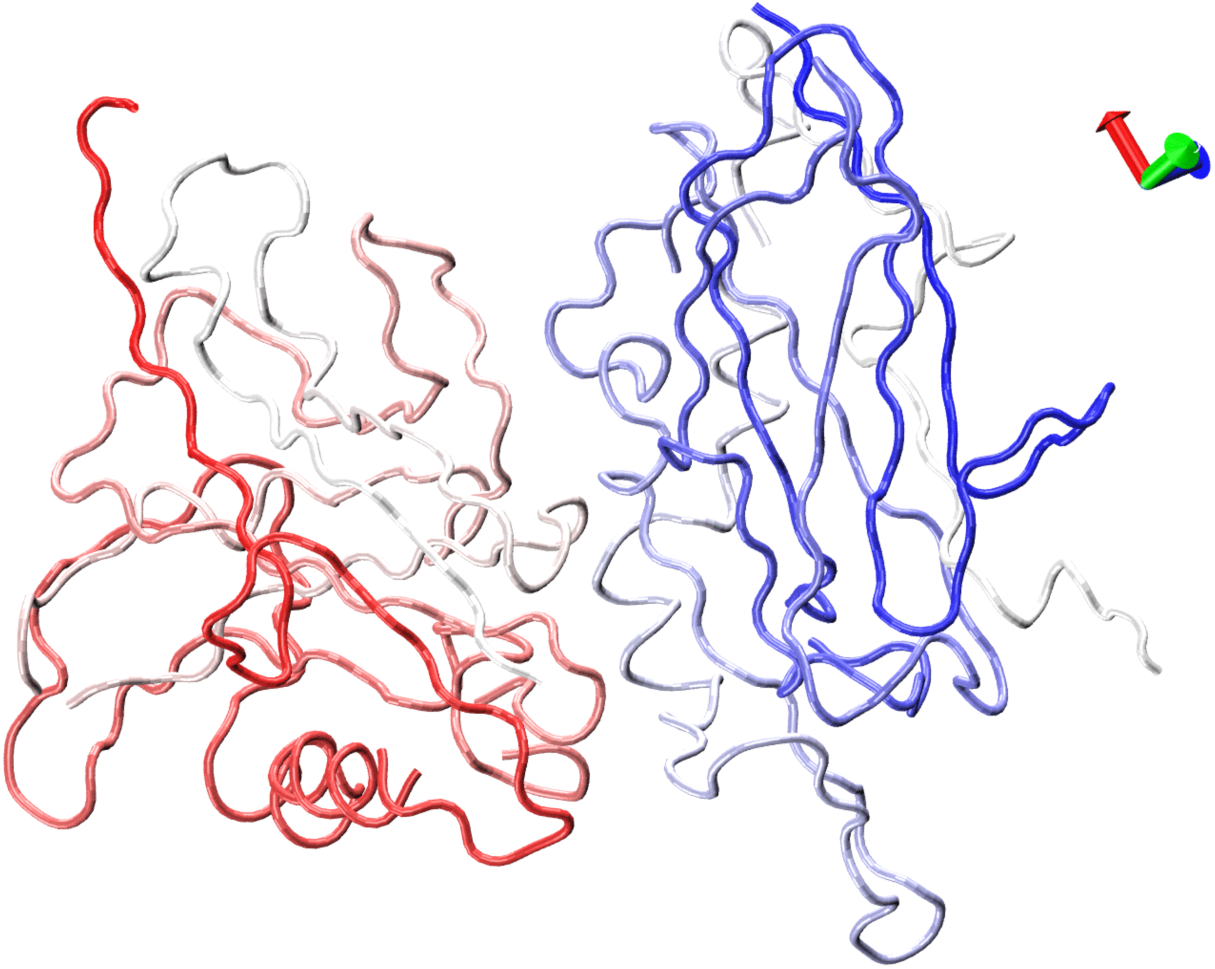
Motion associated with ATP1B2 dimer, PC2.

